# Mutation Rate Variations in the Human Genome are Encoded in DNA Shape

**DOI:** 10.1101/2021.01.15.426837

**Authors:** Zian Liu, Md. Abul Hassan Samee

## Abstract

Single nucleotide mutation rates have critical implications for human evolution and genetic diseases. Accurate modeling of these mutation rates has long remained an open problem since the rates vary substantially across the human genome. A recent model, however, explained much of the variation by considering higher order nucleotide interactions in the local (7-mer) sequence context around mutated nucleotides. Despite this model’s predictive value, we still lack a biophysically-grounded understanding of genome-wide mutation rate variations. DNA shape features are geometric measurements of DNA structural properties, such as helical twist and tilt, and are known to capture information on interactions between neighboring nucleotides within a local context. Motivated by this characteristic of DNA shape features, we used them to model mutation rates in the human genome. The DNA shape feature based models show up to 15% higher accuracy than the current nucleotide sequence-based models and pinpoint DNA structural properties predictive of mutation rates in the human genome. Further analyzing the mutation rates of individual positions of transcription factor (TF) binding sites in the human genome, we found a strong association between DNA shape and the position-specific mutation rates. The trend holds for hundreds of TFs and is even stronger in evolutionarily conserved regions. To our knowledge, this is the first attempt that demonstrates the structural underpinnings of nucleotide mutations in the human genome and lays the groundwork for future studies to incorporate DNA shape information in modeling genetic variations.

## Introduction

The process of genetic mutations bears critical implications for our health and evolution. A nucleotide’s *mutation rate*, defined as its probability of mutating in an individual’s genome, is highly variable in the human and other mammalian genomes (1). Modeling mutation rates in the human genome is therefore a challenging goal. However, predictive modeling of mutation rates and explaining the sources of their genome-wide variations is essential for human genetics. This understanding is the key to studying evolutionary divergence, detecting adaptive evolution (1-3), identifying functional elements in the genome, predicting deleterious variants (1,4-6), and identifying disease subtypes (7-10).

Multiple molecular factors could influence mutation rates, yet none of these factors satisfactorily explains mutation rate variations in the human genome (1,2,11-13). Sequence context, i.e., the DNA sequence flanking a mutated position, has been hypothesized to influence mutation rates strongly. The “CpG context,” for example, explains the 14-fold increase in mutation rates in the context of CpG dinucleotides (1,11,12). Building on this idea, Aggarwala and Voight recently showed that mutation rates estimated from 7-mer sequence contexts (three nucleotides up- and downstream of the mutated positions) could optimally explain mutation rate variations in the human non-coding genome (3). They also showed that these estimates are (a) not influenced by rates of recombination, (b) strongly correlated with rates of species divergence, (c) are consistent for both rare and common genetic variants, and (d) are also reflected in *de novo* mutational events (3). They modeled these estimates using linear regression that incorporated up to fourth-order interactions between nucleotides and explained ∼81% of mutation rate variations in the human non-coding genome (3). Although Aggarwala and Voight’s model is a breakthrough, it is unclear if the higher-order interactions in their model have any biophysical underpinning (3). In this study, we investigate whether it is possible to build a biophysically-grounded yet accurate mutation rate prediction model.

DNA shape features are nucleotide sequence-dependent biophysical features that represent the DNA molecule’s local three-dimensional structures (14-22). DNA shapes are easily interpretable as they represent biophysical properties and essentially capture the interactions between neighboring nucleotides. However, the relevance of DNA shape features has not been systematically explored for mutation rates -- only one study so far has associated DNA curvature with mutation rates in the yeast URA3 gene (23). Taking advantage of population-scale whole-genome sequence data from the 1000 Genomes Project (1KG) (24,25) and the accurate predictions of DNA shape features by the recent DNAShapeR model (19-22), here we explore the relationships between DNA shape features and mutation rate variations in the human genome. We modeled genome-wide mutation rates using DNA shape features and show that DNA shape features can provide biophysically-grounded and up to 15% more accurate models than those with higher order interactions between nucleotide sequence features. Our models reveal that DNA Roll, Shift, Tilt, Stretch, and helical twist (HelT) encode the most information related to mutation rate variations. DNA shapes also characterize mutation promoting genomic contexts with high accuracy (mean auROC, area under receiver operating curve, is 0.9(using a diverse set of predictive features.

Extending the above findings, we finally asked if DNA shape is associated with mutation rate variations in transcription factor binding sites (TFBSs), a large class of potentially functional and evolutionarily selected sequences in the human genome. Indeed, our analysis of 575 transcription factors (TFs) revealed that the mutation rates of different positions of a TF’s binding sites correlate strongly with DNA shape features. The correlations are even stronger in evolutionarily conserved cis-regulatory regions of the human genome. Overall, our study presents the strong relationships between DNA shape and mutation rate variations in the human genome, and lays the groundwork for future genetic studies to incorporate the DNA molecule’s the structural properties.

## Results

### Incorporating DNA shape features improves the performance of mutation rate modeling in the human genome

Aggarwala and Voight recently showed that mutation rates estimated from 7-mer sequence contexts could optimally explain mutation rate variations in the human genome (3). Briefly, for each class of mutation, say A-to-C, they partitioned its occurrences according to the *k* nucleotides flanking the mutated position (⌊*k/2*⌋ nucleotides on both sides). Then, instead of studying the genome-wide A-to-C mutations, Aggarwala and Voight studied *k*_1_-to-*k*_2_ mutations for *k*-mers *k*_1_ and *k*_2_ that differ only in the mutated (middle) position, where *k*_1_ has an A and *k*_2_ has a C. In this setup, *k* = 1 represents the conventional approach for studying mutation rates. However, by increasing the value of *k* from 1 to 7, Aggarwala and Voight showed that the *k*_1_-to-*k*_2_ mutation rates estimated from a training set of human chromosomes correlates significantly better with the mutation rates estimated from a validation set.

Aggarwala and Voight then used these 7-mer sequence context-based mutation rates as the data for modeling mutation rate variation in the human genome. In particular, for a pair *k*_1_ and *k*_2_ of 7-mers that differ at the central (mutated) position, they modeled *k*_1_ *→k*_2_ mutation rates from the nucleotides in *k*_1_ and *k*_2_. They found a linear regression model with up to fourth-order interaction terms to be optimal for this data, explaining ∼81% of the variance in mutation rates (3). Although this performance is remarkable, there remains open questions on (a) do the nucleotide interactions in their models carry any biophysical significance? and (b) could a more biophysically grounded model improve this performance? Since DNA shape is a biophysical representation of interactions between neighboring nucleotides (18), we posited that modeling genome-wide mutation rates using DNA shape features could offer biophysical insights without reducing the model’s performance. We pursued this goal using a series of increasingly complex models from DNA nucleotide and shape features, and selected the optimal models using cross-validation strategy.

Like Aggarwala and Voight, we separately modeled nine mutation classes after combining mutation rates from reverse-complement sequences (Figure 1A). If the mutated nucleotide was a cytosine (C), Aggarwala and Voight had considered two mutation classes based on whether a guanine (G) follows the cytosine (CpG context) or not (non-CpG). We fit the models with nucleotide and shape features (Figure 1B-C), either alone or in combination, while systematically increasing model complexities by including higher-order interactions (Figure 1C). Following Aggarwala and Voight, we allowed for up to fourth-order interactions between nucleotide features. Following previous works using DNA shape for modeling the DNA-binding specificity of TFs, we also allowed second-order interactions between DNA shape features from adjacent positions (19,27,28). We trained each model by optimizing L1-penalized mean squared error loss, and selected the optimal model based on 8-fold cross validation (Figure S1, Methods). We then tested each model on a separately held-out test data and confirmed the model performances are consistent between training and testing data (Figure S1, Table S1-2).

**Figure 1.**
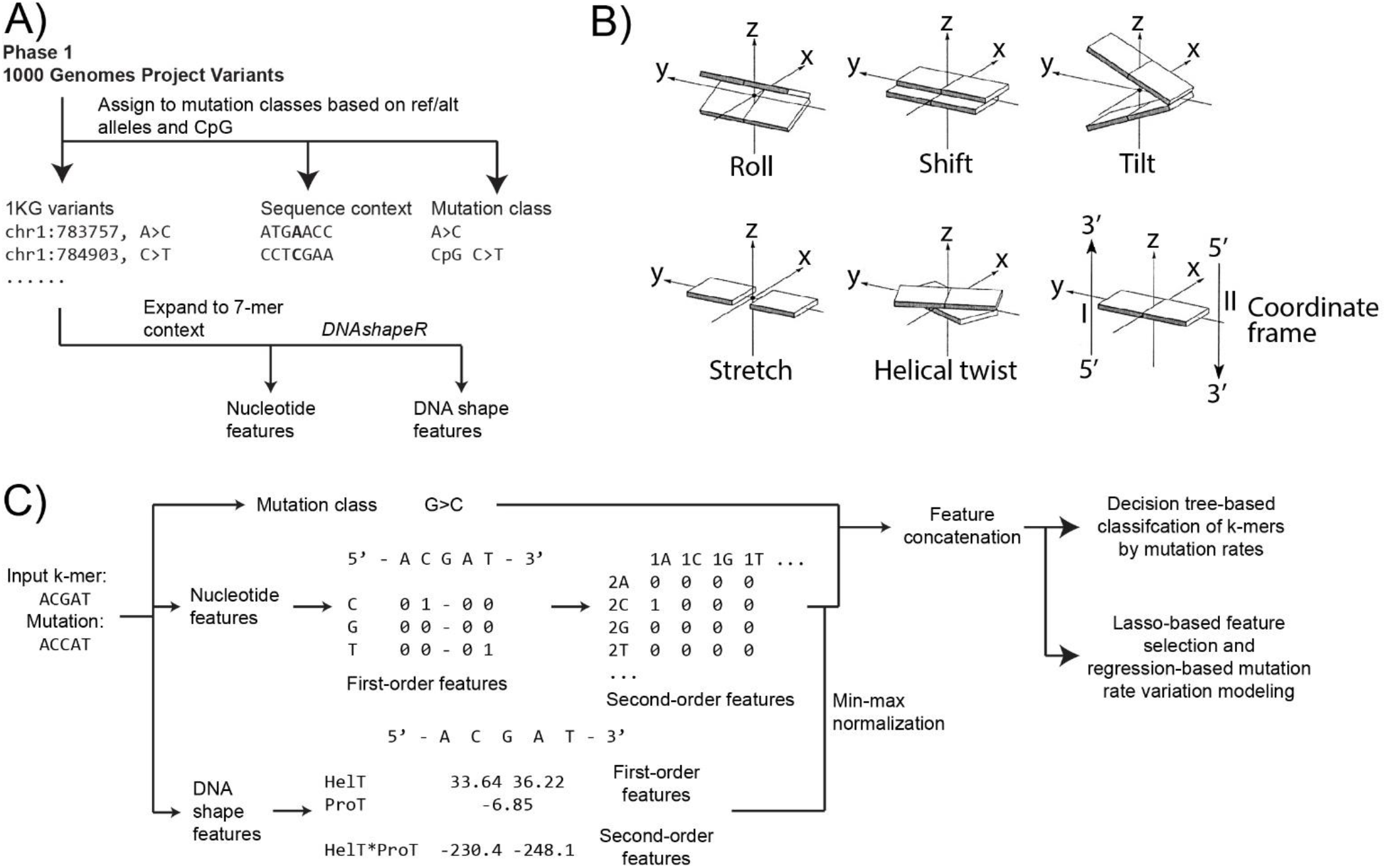
Overview of our study. A) Our pipeline for the generation of predictive features from mutation data. Mutation data were obtained from phase 1 of the 1000 Genomes Project (1KG) as described in the Methods section. Mutations were then extended into 7-mers based on the identities of the three flanking nucleotides; within each 7-mer context, mutation rate will be calculated by dividing the total number of mutations with that particular 7-mer context with the total number of the corresponding 7-mer context observed in the human noncoding genome. B) Illustrations of several first-order DNA shape features as used in our study, including Roll, Shift, Tilt, Stretch, Helical twist (HelT), as well as an illustration of the axes. Data were obtained from 3DNA (26). C) Example of our machine learning pipeline given an input 5-mer of ACGAT, note that we listed a 5-mer example here for simplicity, but we only used 7-mers in the study. For nucleotide features, we will generate three corresponding first-order features corresponding to C, G, and T for each location on the k-mer; we have excluded the central location as the central location will become a uniform feature due to us modeling each mutation class separately. For shape features, the *DNAshape* method generates 14 unique types of shapes, for which we show two examples (helical twist, HelT; propeller twist, ProT) here. Since HelT is an inter-basepair feature and ProT is an intra-basepair feature, there are two HelT and one ProT values for each 5-mer context, see the methods section for more information. We also show one example of interaction feature HelT × ProT, for which we will assume to be inter-basepair if any one of the two interacting features is inter-basepair.

Across all nine mutation classes, the models containing second-order shape interactions or combining shape and nucleotide features outperformed the current best model of Aggarwala and Voight (Figure 2A-C, Table 1, Figure S2-3, Table S1-S2). The models showed a 3% improvement in median *R*^2^, with the highest improvement being 15% in C-to-A mutations in CpG context. The other classes that benefitted the most from shape features are A-to-T, C-to-A (both CpG and non-CpG), and C-to-G. Below we further tease out the interplay between nucleotide and shape features and interpret our models in terms of DNA shape.

**Table 1.** Model performance of our best-performing model in the testing data. The “maximum achievable R^2^” column values were calculated by directly comparing testing data with the training data and represent “maximum” model performances. The “Aggarwala” column describes performances of the current state of the art model. Performance percentage changes were calculated by 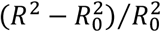 comparing our model to the state of the art model. The “parameters” column describes the identities of the input predictors for each mutation class. Regarding the abbreviations in the parameters column, numbers represent degrees of polynomial transformations, “sh” and “sc” represent shape and nucleotide (sequence context) features, and the “neibr” suffix represents that only neighboring interactions were included (see Methods). The mean and median values were computed based on R^2^ values of the nine individual models and were not computed for performance changes to avoid ambiguity.

| Mutation class | Maximum achievable $R^2$ | Aggarwala | Liu | Performance change | Parameters |
| --- | --- | --- | --- | --- | --- |
| A>C | 0.666 | 0.580 | 0.592 | 1.96% | sh2neibr_sc4 |
| A>G | 0.961 | 0.920 | 0.931 | 1.24% | sh2neibr_sc4 |
| A>T | 0.750 | 0.605 | 0.673 | **11.18%** | sh2neibr |
| C>A | 0.905 | 0.840 | 0.882 | 5.01% | sh2neibr_sc4 |
| C>G | 0.867 | 0.819 | 0.844 | 3.03% | sh2neibr_sc1 |
| C>T | 0.864 | 0.875 | 0.882 | 0.77% | sh2neibr_sc4 |
| CpG_C>A | 0.534 | 0.553 | 0.635 | **14.68%** | sh2neibr_sc3 |
| CpG_C>G | 0.417 | 0.531 | 0.530 | -0.07% | sh2neibr_sc4 |
| CpG_C>T | 0.934 | 0.932 | 0.934 | -0.20% | sh2neibr_sc4 |
| **Mean** | 0.766 | 0.740 | 0.767 |  |  |
| **Median** | 0.864 | 0.819 | 0.844 |  |  |

**Figure 2.**
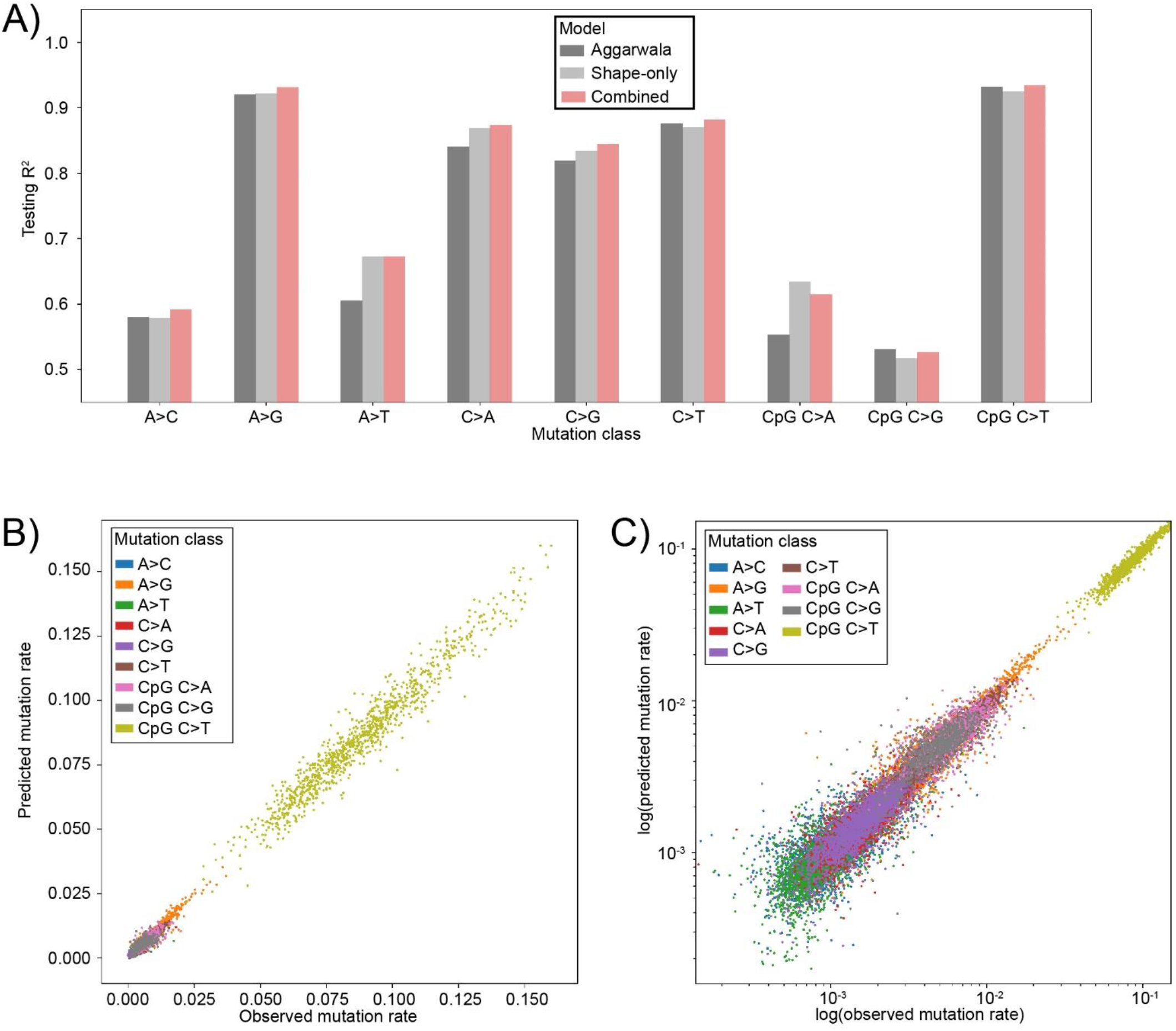
Our best performing model outperforms the Aggarwala and Voight model. A) Histogram comparison of R^2^ values of the Aggarwala and Voight model (*Aggarwala*) (3), our *shape-only* model, as well as our *combined* model on the independent testing data. B-C) Scatterplots showing comparison of predicted mutation rates and observed mutation rates from the 1KG data, in B) linear scale and C) logarithmic scale. The x-axes show observed mutation rates, while the y-axes show model-predicted mutation rates.

### DNA shape features alone, with at most second-order interactions, improve over the current best models that use nucleotide features and fourth-order interactions

Given the above improvements in model performance through introducing DNA shape features, we asked whether and to what extent shape features could replace nucleotide features. Thus, we focused on the models that use DNA shape features alone. As noted above, these models were allowed at most second-order interactions between shape features, and the interactions were limited to feature values from adjacent positions. In the following, we refer to these models as *shape-only* models. We use the term *sequence-only* models to refer to Aggarwala and Voight’s models utilizing nucleotide features with up to fourth-order interactions, and the term *combined* models to refer to the models described above that use both sequence and shape features.

Importantly, our shape-only models showed a 1.8% improvement in median *R*^2^ over the sequence-only models (Table 2, Figure S4-5, Table S2), with the largest improvements in mutation classes A-to-T (11%) and CpG C-to-A (15%), a 3.3% improvement in the class C-to-A, a 2.5% drop in the class CpG C-to-G, and < 1% change in three other classes (Table 1-2, Table S2).

**Table 2.**
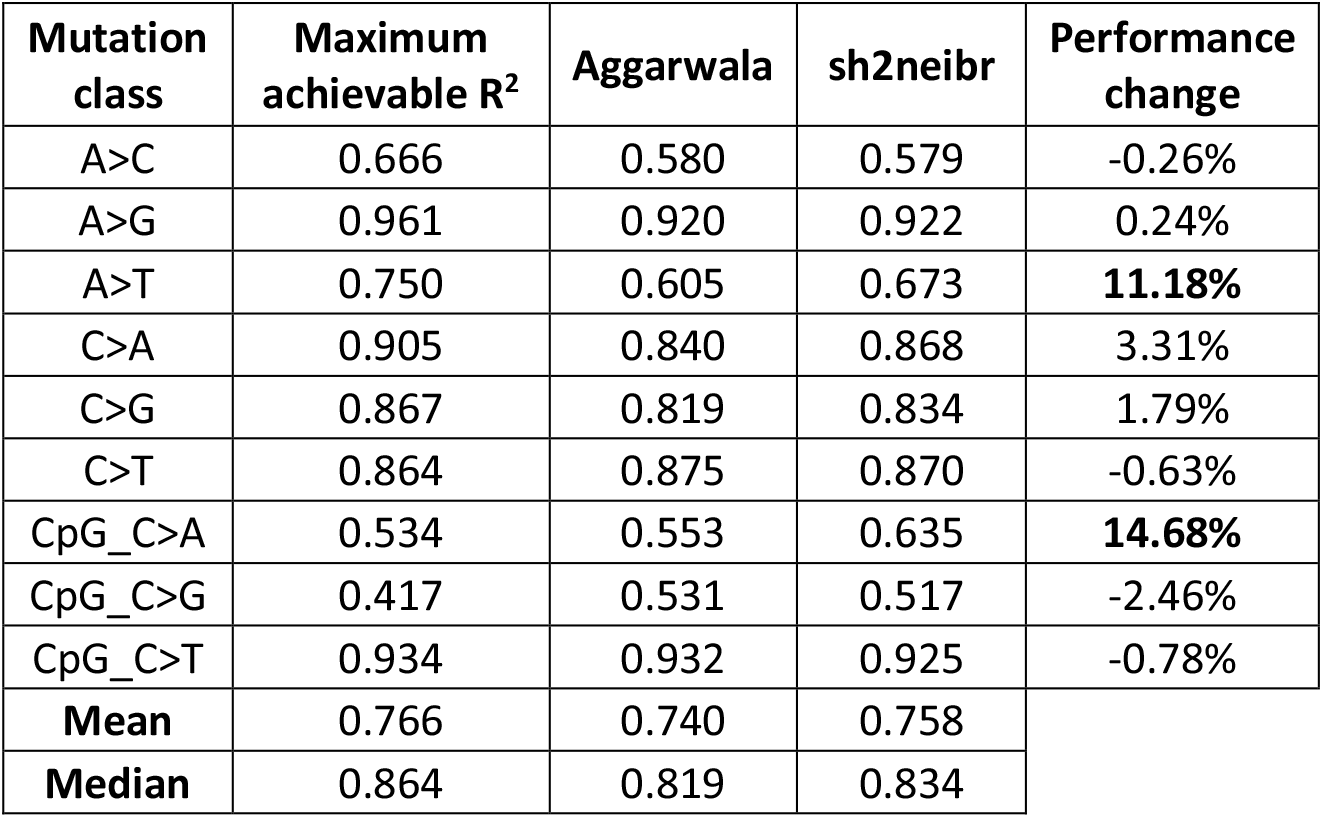
Model performance of our second-order shape model in the testing data. The “maximum achievable R^2^” column values were calculated by directly comparing testing data with the training data and represent “maximum” model performances. The “Aggarwala” column describes performances of the current state of the art model. The “sh2neibr” column describes performances of our second-order DNA shape model. Performance percentage changes were calculated by 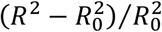 comparing our model to the state of the art model. The mean and median values were computed based on R^2^ values of the nine individual models and were not computed for performance changes to avoid ambiguity.

| Mutation class | Maximum achievable $R^2$ | Aggarwala | sh2neibr | Performance change |
| --- | --- | --- | --- | --- |
| A>C | 0.666 | 0.580 | 0.579 | -0.26% |
| A>G | 0.961 | 0.920 | 0.922 | 0.24% |
| A>T | 0.750 | 0.605 | 0.673 | **11.18%** |
| C>A | 0.905 | 0.840 | 0.868 | 3.31% |
| C>G | 0.867 | 0.819 | 0.834 | 1.79% |
| C>T | 0.864 | 0.875 | 0.870 | -0.63% |
| CpG_C>A | 0.534 | 0.553 | 0.635 | **14.68%** |
| CpG_C>G | 0.417 | 0.531 | 0.517 | -2.46% |
| CpG_C>T | 0.934 | 0.932 | 0.925 | -0.78% |
| Mean | 0.766 | 0.740 | 0.758 |  |
| Median | 0.864 | 0.819 | 0.834 |  |

### Structural underpinnings of mutation rate variations in the human genome

We next analyzed our shape-only models to interpret mutation rate variations from a structural perspective. Since we incorporate up to second-order interactions and select variables through a Lasso regularization, a shape feature could appear alone or with another feature in a second-order variable in these models. We thus defined a shape feature’s *utilization* in a model as how frequently it appears in the model’s variables (Fig. 3A). We also normalized the variables’ coefficients to make them comparable across mutation classes and defined a variable’s *importance* in a model as the absolute value of its normalized coefficient (Fig. 3A). Model coefficients were documented for our *combined* and *shape-only* models (Table S3-4).

**Figure 3.**
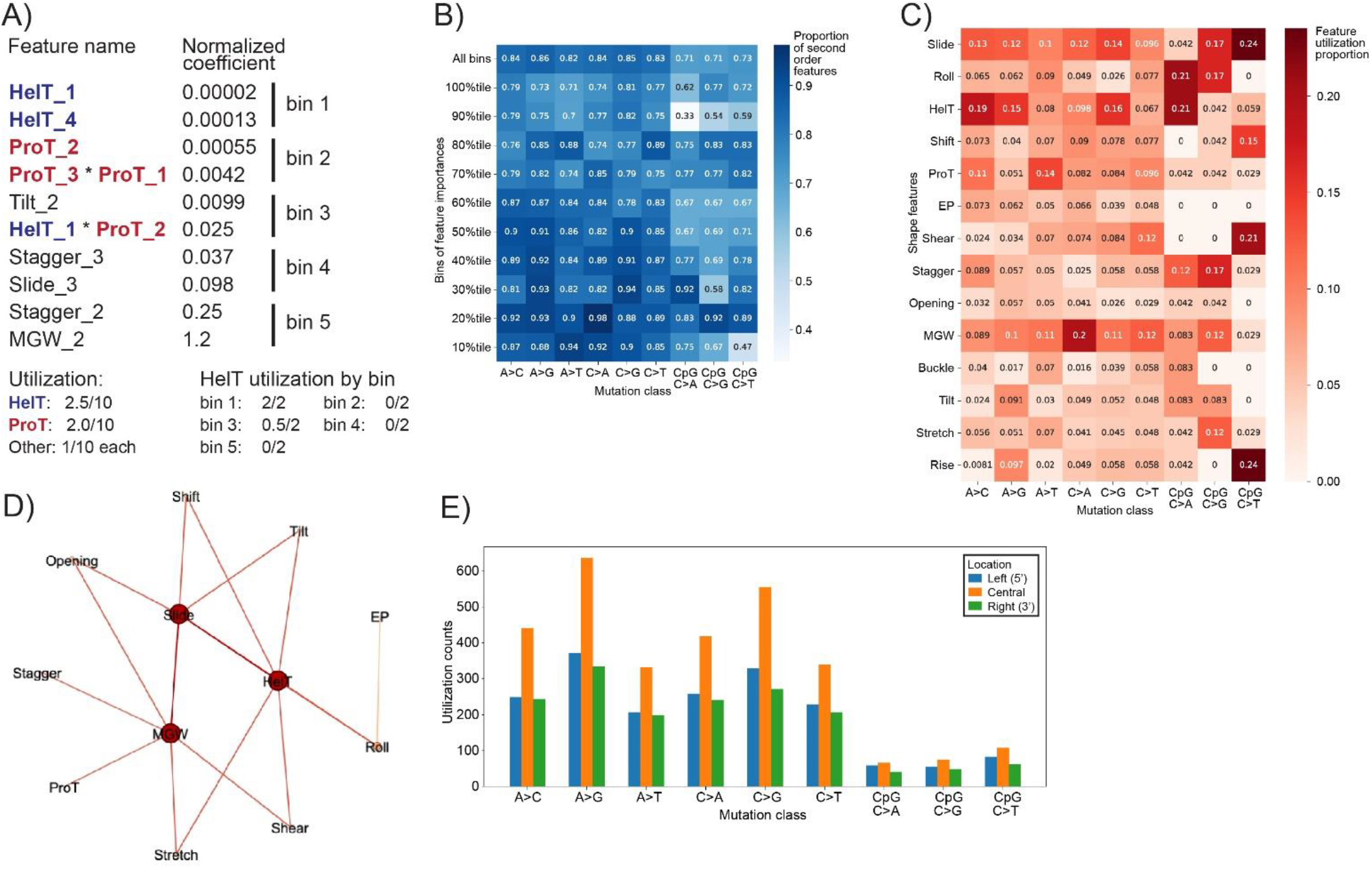
Coefficient analyses of DNA shape-based regression model suggest feature utilization trends. A) Illustration of our calculation of feature utilization using a toy example with 10 features, of which two are interacting features. B) Ratio of second-order shape features among all features used in each of the nine mutation classes as well as in 10% bin increments of importance. The first row describes ratios in the entire models, while the subsequent rows describe ratios in each bin of importance. See Methods for how the bins are defined. C) Relative utilization of 14 shape features across models in all nine mutation classes using top 10% features with the highest importance values. D) Illustration showing pairs of features included in at least 1% of the 10% features with the highest importance values. E) Bar charts of feature utilization based on feature location across models in all nine mutation classes. Feature utilization scores were calculated by counting the number of times a feature is used, and relative feature utilization scores were calculated by dividing the feature utilization scores by the total number of features in a particular model or a particular subset of features.

We noticed that Roll, Shift, Tilt, and Stretch are consistently the most utilized features across all nine mutation classes (utilization of ∼10%, Figure S6). The features ProT (propeller twist), Shear, Stagger, MGW, and Buckle had utilizations between 5% to 10%; and the least utilized (<5%) features were HelT, EP, Opening, and Rise (Figure S6). Interestingly, HelT is highly utilized in the most important (top 10%) variables despite being overall underutilized (Figure 3C, Table 3, Table S5). In contrast, the most utilized features like Roll, Shift and Tilt are common in the less important variables (Methods; Table S5).

**Table 3.** Fisher’s exact tests with Benjamin-Hochberg false discovery rate (FDR-BH)-adjusted p-values of feature utilization between top 10% features and all features. The top 10% features were defined based on feature importance, or absolute coefficient values.

| Shape features | A>C | A>G | A>T | C>A | C>G | C>T | CpG C>A | CpG C>G | CpG C>T |
| --- | --- | --- | --- | --- | --- | --- | --- | --- | --- |
| Slide | 0.152 | 0.106 | 0.468 | 0.297 | 0.166 | 0.839 | 1 | 1 | 0.220 |
| Roll | 1 | 1 | 1 | 1 | 1 | 1 | 1 | 1 | 1 |
| HelT | **0.000349** | **0.00132** | 0.498 | 0.114 | **0.00132** | 0.857 | 0.498 | 1 | 1 |
| Shift | 1 | 1 | 1 | 1 | 1 | 1 | 1 | 1 | 1 |
| ProT | 1 | 1 | 0.498 | 1 | 1 | 1 | 1 | 1 | 1 |
| EP | 1 | 1 | 1 | 1 | 1 | 1 | 1 | 1 | 1 |
| Shear | 1 | 1 | 1 | 1 | 1 | 0.857 | 1 | 1 | 0.650 |
| Stagger | 1 | 1 | 1 | 1 | 1 | 1 | 1 | 1 | 1 |
| Opening | 1 | 1 | 1 | 1 | 1 | 1 | 1 | 1 | 1 |
| MGW | 1 | 0.971 | 0.860 | 0.021 | 0.857 | 0.583 | 1 | 1 | 1 |
| Buckle | 1 | 1 | 1 | 1 | 1 | 1 | 1 | 1 | 1 |
| Tilt | 1 | 1 | 1 | 1 | 1 | 1 | 1 | 1 | 1 |
| Stretch | 1 | 1 | 1 | 1 | 1 | 1 | 1 | 1 | 1 |
| Rise | 1 | **0.0384** | 1 | 0.498 | 0.545 | 0.766 | 1 | 1 | **0.0384** |

Further analyzing the most important variables in each mutation class, we found that the majority (>90%) of these variables are second-order (Table S4). These generally represent the interaction between two different shape features, although there was a high variation in this aspect across the mutation classes. For example, in the C-to-T mutation model within the CpG context, 47% second-order variables represent interaction between two different shape features, but this fraction is 85% in the C-to-T mutation model outside the CpG context. These second-order variables also revealed that HelT, Slide, and MGW interact most frequently with other shape features. Interestingly, whereas the base-step features commonly interact with each other, the base-pair features show interactions mostly with MGW but not within themselves (Figure 3D).

Since a DNA shape feature’s values change depending on its location in the 7-mer, we then asked if the variables selected in our models include shape feature values uniformly from all locations. We found that shape features around the central (mutated) position are more utilized than those in the adjacent positions (Figure 3E). Supplemental figure).

### DNA shape can accurately identify mutation-promoting sequence loci

The 7-mer context-based mutation rates could vary by up to 10-folds between the most and the least frequently mutated contexts (Figure S8). Distinguishing the most frequently mutated 7-mer contexts can elucidate the mutation-promoting and DNA repair characteristics of the genome (3), but a regression model may not necessarily reveal this information. Thus, to explore whether and how DNA shape may cause certain sequence contexts to have very high or very low mutation rates, we built classification models to distinguish between sequence contexts showing the highest and the lowest 5 and 10% mutation rates. Using simple two-layer decision trees with first-order shape features, we obtained highly accurate classification models (mean area under receiver operating curve, auROC = 0.9) that discriminates sequence contexts showing the highest and the lowest 5% mutation rates. The models’ auROCs were equally high for the highest and the lowest 10% mutation rates (Figure 4A, Figure S9, Table S6-8).

**Figure 4.**
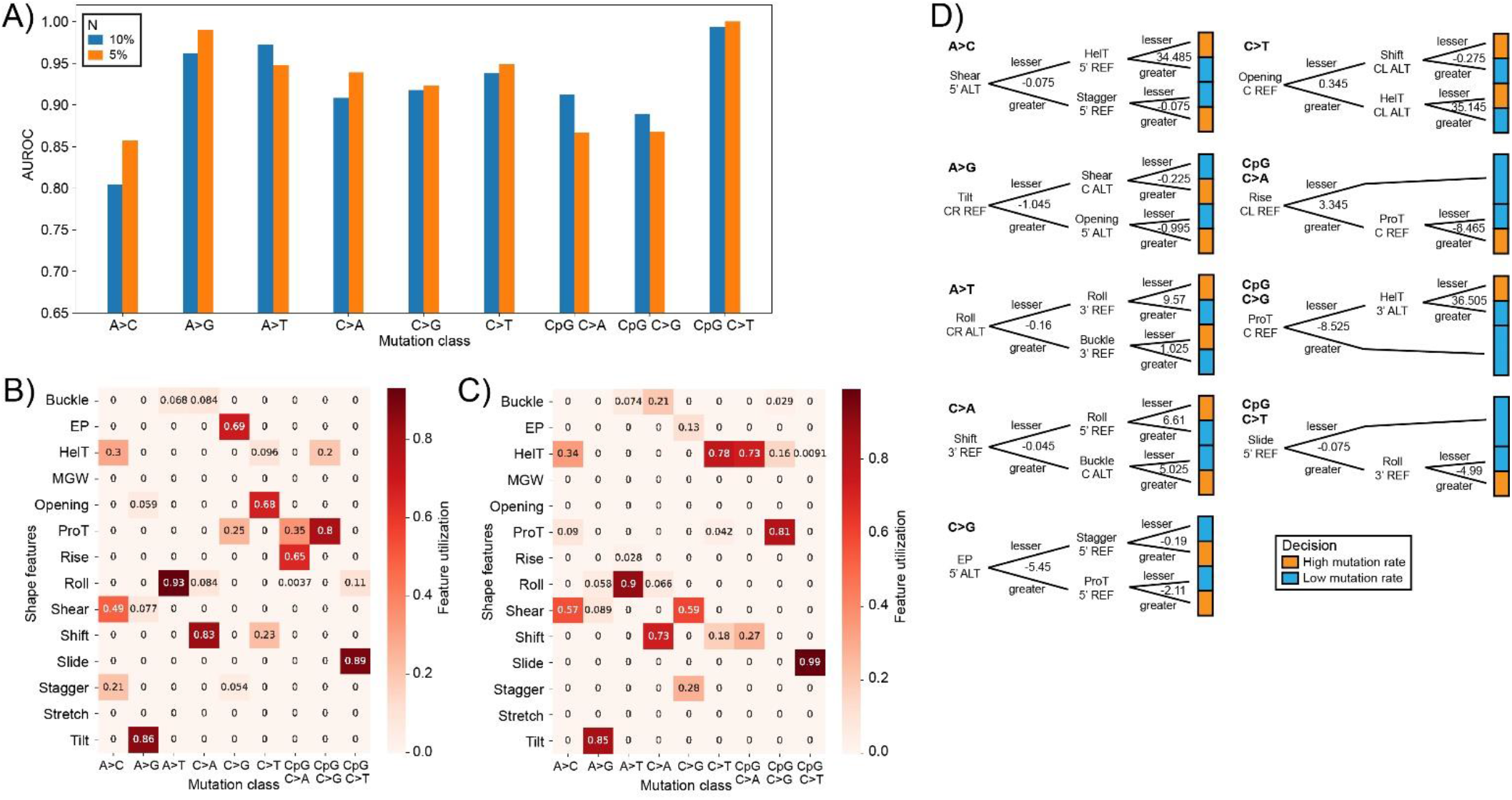
Decision tree-based classification models show high accuracy and interpretability for classifying k-mers with the highest or lowest mutation rates. A) Bar chart of area under receiver-operating curve (AUROC) values for classification models for all nine mutation classes and values of N equal to 5% or 10%. The ROC curves were constructed using false positive rates and true positive rates. B-C) Heatmaps of feature importance, as calculated by the total reduction in Gini coefficient by a particular feature, across all shape features (y-axis), all nine mutation classes (x-axis), and values of N equal to B) 5% and C) 10%. See Methods section for details on feature importance calculations. D) Detailed decision tree architectures for N=5 in all nine mutation classes. Each decision tree feature is labeled by the type of shape feature, location, and strand; possible locations include 5’, central-left (CL) and central-right (CL) for inter-basepair features, central (C) for intra-basepair features, and 3’; possible strands include reference (REF) and alternative (ALT). Note that for all subplots, N represents the percentage of k-mers with the highest or lowest mutation rates and were used as inputs to the classification models, see the methods section for more information.

Interestingly, unlike the regression models, the most important shape features in the classification models corresponded to completely different shape features across the nine mutation classes (Figure 4B-C; based on Gini index; see Methods). In most cases, the most important shape features are base-step features (Figure 4B-D) and in a few classes, such as in A-to-G, A-to-T, and C-to-T in the CpG context, the root feature (feature selected in the first layer of the decision tree) alone could make highly accurate predictions (Figure 4B-D, Figure S10). Features from all locations in the 7-mer, as well as both reference and alternative 7-mers, were roughly equally represented across the root features (Figure 4B-D, Figure S10).

For orthogonal validation, we classified all 7-mers that fall into extended mutation promoting or inhibiting sequence motifs as characterized by Aggarwala and Voight (3). We have first shown that across all possible 7-mers belonging to these motifs, our models have reached very high agreements with the Aggarwala and Voight results (Table S9). We have then selected six exemplar 7-mers belonging to those sequence motifs as well as the shape features that our models used to reach such conclusions (Table S10); note that our models have reached consensus on the classification of all six 7-mers.

Overall, this objective analysis showed DNA shape can characterize the mutation promoting and mutation averting sequence contexts with high accuracy. To our knowledge this is the first such thorough characterization of mutation promoting and averting sequence contexts; the previous analysis in this realm was based on enrichment in the highest and the lowest 1% mutation rates (3). However, the overall classification accuracy achievable through those enriched sequences was unclear. It was also unclear if it would be possible to characterize a larger set of sequences (5% or 10%) with such high accuracy.

### DNA shape alterations correlate negatively with mutation rates in transcription factor binding sites

Extending the above relationship between DNA shape and genome-wide mutation rates, we finally asked if a similar relationship exists in potentially functional genomic regions. To this end, we focused on the mutation rates of individual positions of TFBSs (29). Previous works have termed this statistic as “per position diversity across individuals” and “variation frequency at motif positions” (29,30).

We analyzed Kheradpour and Kellis’ curated list of genome-wide TFBSs for 575 TFs (median of ∼53,000 sites per TF motif) (31). TFBSs are known to be less polymorphic in general (32), but we still noted the mutation rates across the positions of a TF’s binding sites could vary by a median of 3-folds (Figure S11). To relate the mutation rates of each position of a TF’s binding sites with the resulting alterations in a DNA shape feature’s values, we first derived the shape feature’s distributions from the reference and the alternate alleles and computed the Kolmogorov-Smirnov (KS) statistic between these distributions. We then compared the KS statistic of each position with the mutation rate at that position. As we show for the example of CTCF (CCCTC-binding factor), this analysis revealed a consistent signal of negative correlation between DNA shape changes and mutation rates (Figure 5A-B). This anti-correlation trend holds for the vast majority TFs (Fig 5C-D), with the median Spearman correlation between the KS statistic of shape features and mutation rates being −0.2 (Figure 5D, Table S11). The anti-correlation values are strongest on average for Homeodomain, bHLH (basic helix-loop-helix), and Tryptophan TFs (Figure 5C).

**Figure 5.**
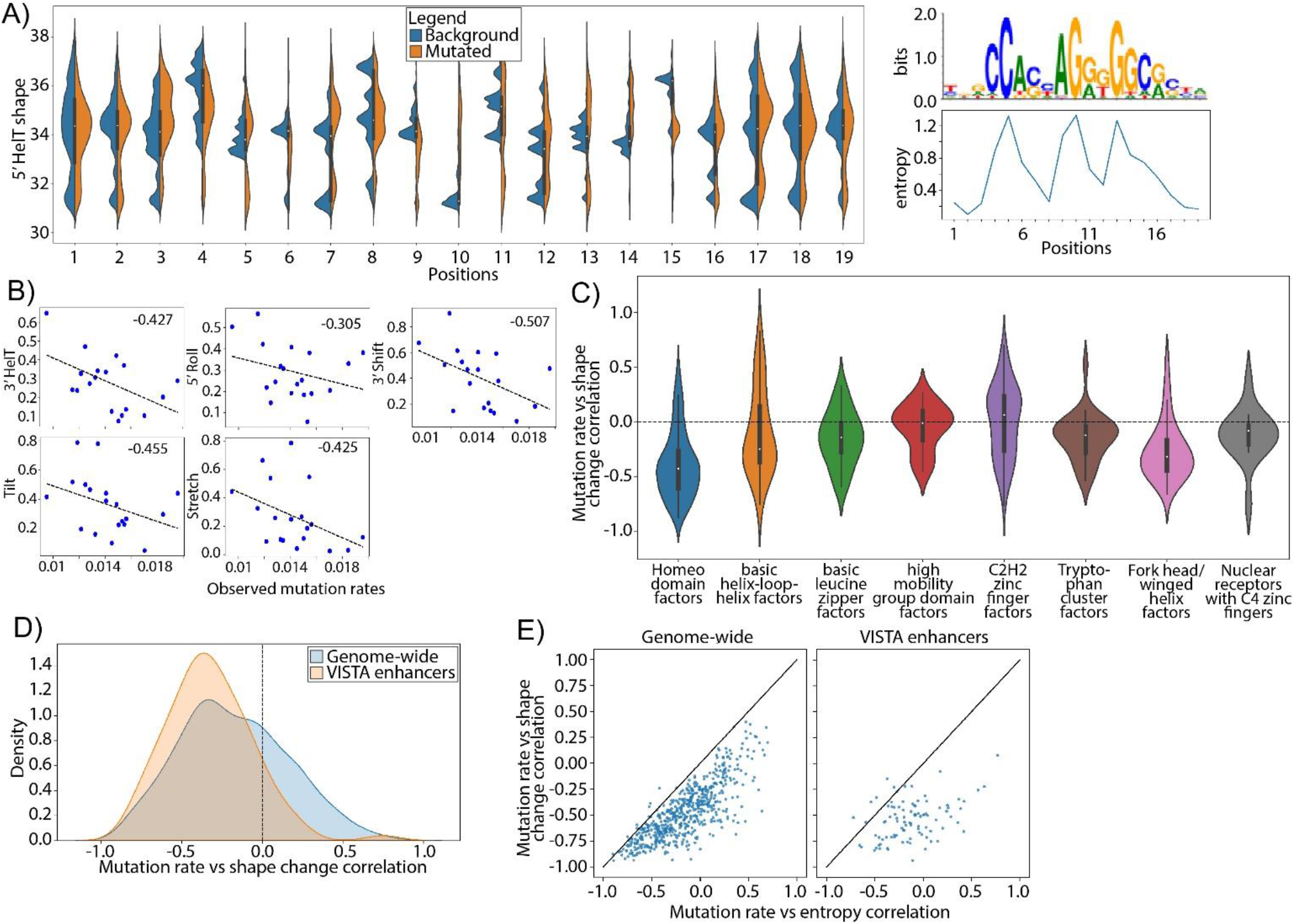
Mutations are less likely to occur at positions that would cause significant shape deviations from native DNA shape distributions. A) Sequence logo and entropy values (right), as well as paired violin plots of the distributions of helical twist shapes (left) of the *CTCF_known1* motif. B) Scatterplots showing correlations between KS statistics of shape changes of HelT-3’ (top-left), Roll-5’ (top-middle), Shift-3’ (top-right), Tilt (bottom-left), and Stretch (bottom-middle) versus per-position mutation rates across the CTCF_known1 motif. Annotated values show spearman rank correlation values between the two variables, note that Shift-3’ has the strongest correlation with per-position mutation rates across all shape features. C) Violin plot of distribution of correlation values between mutation rates and KS statistics of shape changes by TF family designations. Only families with more than 10 TFs in our dataset are included, and TFs in the “unspecified” family are not shown. D) Distribution of spearman rank correlation values between observed mutation rates and KS statistics of helical twist shape changes for all transcription factors and transcription factors that commonly occur in VISTA enhancer regions. E) Scatterplots showing paired correlation values for each TF among all transcription factors or those that commonly occur in VISTA enhancer regions. The x-axes denote correlation values between mutation rates and entropy, and the y-axes denote correlation values between mutation rates and KS statistics of shape changes, for which the shape feature with the most negative correlations were used.

We note that the above finding aligns with Spivakov et al.’s analysis of evolutionarily conserved binding sites for 36 human TFs (30) where information content per motif position and mutation rates showed a median anti-correlation of −0.2. Since DNA shape data is derived from sequence data, we expected such confirmatory findings. Indeed, for all TFs, we found the above KS statistic is strongly correlated with information contents per motif position (Figure S12). However, our analysis suggests the association between DNA shape alteration and mutation rates on a larger scale (for 575 TFs) and genome-wide beyond evolutionarily conserved sites. Importantly, the anti-correlation signal between shape alteration and mutation rates becomes even stronger (−0.5) when we focused on evolutionarily conserved regulatory regions in the human genome (Figure 5D-E, Table S12). Overall, in the context of mutation rates, we find that DNA shape captures a signal complementary to nucleotide sequences.

## Discussion

We have shown here the efficacy of DNA shape in explaining the genome-wide single nucleotide mutation rate variations in the human genome. Our models not only outcompeted the current best models, but also offered novel insights into the structural underpinnings of single-nucleotide mutations. We have also built simple decision tree-based models to categorize the sequences that are more likely to mutate, and used our models to interpret several mutation-promoting or inhibiting motifs described by Aggarwala and Voight (3). In addition, we have also taken our findings to the context of TFBS and discovered that mutations that significantly perturb DNA shapes are less likely to occur in functional genomic regions.

To our knowledge, this the first detailed investigation relating DNA shape and single nucleotide mutation rates. A previous work has commented on DNA curvature having a negative correlation with mutation rate in the *URA3* gene (23). Two features of DNA curvature, Roll and Tilt, were moderately utilized across all mutation classes except for CpG C-to-T, and appeared as root features for decision tree models for the mutation classes A-to-G and A-to-T. This is an encouraging piece of evidence that our finding has echoed with previous literature.

Our models revealed two critical points on the role of DNA shape and the necessity of considering the exact nucleotide’s identity in modeling mutation rates. First, we found that using only DNA shape and without any nucleotide sequence features, we were able to outperform the current base sequence feature-based models. Furthermore, when we interpreted the models, we found similar features to be the most important and the most utilized for all mutation classes. This demonstrates that DNA shape features encode essential information to explain mutation rate variations. On the other hand, like the previous works in this realm, we needed to limit our modeling to individual mutation classes. This necessity to maintain the information of the mutated nucleotides’ identities implies that there could be additional shape features or other properties of the DNA molecule that we still lack in the models.

We have excluded non-neighboring DNA shape interactions for all our reported models. It is worth noting that we have compared the performances of models that include or exclude non-neighboring shape interactions. Removing non-neighboring interactions in DNA shape features had little adverse effect on model performance (Table S13-14) but reduces the number of input features from 4158 to 2089 (see Methods). This implies that the neighboring shape interactions capture all necessary information for predicting mutation rate variations, thus prompting us to remove non-neighboring shape interactions.

Ideally, we would expect that going beyond 7-mers would capture more shape-specific information and improve the models further. Although we have attempted using up to 9-mer local sequence context to explain more variance in mutation rates, we found many 9-mer mutation patterns were not observed within the Phase 1 1KG and resulted in data sparsity (24). We also note that at the level of k >= 11, “nullomers”, DNA k-mer sequences expected to occur in the human genome but did not, will occur (33-35). This leads us to a hypothetical maximum k of 9 for our local sequence context models should we expand our choice of k to increase the amount of variance explained. It has been well established by literature that CpG sites significantly elevate mutation rates (1,11-13). It is also known that 70 − 80% of all CpG sites in adult human tissue are permanently methylated with another ∼20% showing dynamically methylation (36). As such, we built our models under the assumption all instances of CpGs in our input 7-mers are methylated. Hence, we have used methylated shapes in DNAshapeR for the four shapes (Roll, ProT, HelT, MGW) that have it available (see Methods). Interestingly, HelT proved to be one of the most important features in both our regression and classification models (Figure 3C, Figure 4B-C, Table 3). In the future, as the methylated values of more DNA shape features become available, we will update our models to better capture the dynamic DNA methylation landscape.

The above analyses established the suitability of DNA shape features in modeling genome-wide mutation rate variations in an accurate and interpretable manner. We finally asked if, in a reciprocal manner, these mutation rate models have provided us with any novel insight on DNA shape. As discussed above, Aggarwala and Voight’s sequence-only model allows nucleotide interactions between any combination of up to four nucleotides within the 7-mer context of a mutated position (3). However, for DNA shape features, even when we limited their interactions to only adjacent positions, the shape-only models with up to second-order interactions were consistently as good as the sequence-only models (Table S13-14). This difference between the two models has critical mechanistic implications on the ability of DNA shape in capturing the effects of nucleotide interactions on mutation rates. It has been suggested that DNA shape features essentially capture the interactions between di-nucleotides, i.e., second-order interactions between neighboring positions (37). This implies that a sequence-only model with up to fourth-order nucleotide interactions, but the interactions limited to only adjacent positions, would be able to capture the effects of DNA shape. Thus, limiting nucleotide interactions to adjacent positions should not lower the performance of Aggarwala and Voight’s sequence-only model compared to our shape-only model. A contrary would suggest that some DNA shape features capture nucleotide interactions between non-neighboring nucleotides.

We found that limiting nucleotide interactions to only neighboring positions caused up to 23% drops in explained variance of the sequence-only models across different mutation classes (Table S14). This suggests that some DNA shape features indeed capture nucleotide interactions beyond non-neighboring nucleotides. To identify the cases where DNA shape features capture nucleotide interactions beyond di-nucleotides, we asked if combining DNA shape features with the above sequence-only model where interactions are limited to adjacent positions could raise the model’s performance back to Aggarwala and Voight’s sequence-only model. Incorporating first-order DNA shape features to above model indeed recovered the performance to a large extent for all classes (Table S13-14). Furthermore, incorporating second-order shape features almost entirely recovered the performances, although we limited the shape features’ interactions to only neighboring positions (Table S13-14). The notable exceptions here were C-to-G mutations (both within and outside the CpG dinucleotide context). Overall, this analysis suggested that some DNA shape features capture nucleotide interactions beyond di-nucleotides and second-order DNA shape feature interactions, even if limited to adjacent positions, are generally sufficient to capture complex higher order nucleotide interactions.

With increasing improvements in population-scale human variation data, such as the Genome Aggregation Database (38), and predictive modeling to derive the structure of arbitrary DNA sequences (18-22,39), our study opens up new possibilities to mechanistically understand mutation rate variations in the human genome. In future studies, we may be able to use broader (9-mer) local contexts for predicting mutation rates, which we failed to do in our current study due to limitations from the 1000 Genomes dataset.

## Methods

### Mutation rate data

We used Aggarwala and Voight’s estimates of single nucleotide mutation rates in the human genome (3). Considering the 7-mer context around each mutated nucleotide, they estimated the rates from the African population in Phase 1 of 1KG (24). Briefly, they first filtered the 1KG variants by population and excluded all annotated genes, centromeres, telomeres, repetitive regions, and regions not annotated in the accessibility mask of 1KG. Then, they calculated the count of each 7-mer, as well as the count of each unique mutation pattern along with their 7-mer context, separately for each human autosome. Finally, they computed the training and test data of mutation rates by combining the counts from all even-numbered autosomes (training data) and from all odd-numbered autosomes (testing data). See Figure 1A for an overview of the steps and Figure S1 for an overview of how we modeled the data.

### DNA shape data

We used the R package DNAshapeR to obtain DNA shape features of the mutated nucleotides and their flanking sequences (19-22). DNAshapeR provides DNA shape data of 14 physio-chemical features estimated from all-atom Monte Carlo simulations (19-22). Given an input DNA sequence, DNAshapeR scans the sequence in a sliding window of length five and outputs the DNA shape features of each 5-mer window in the sequence. The 14 features classify into three types: inter-base pair shapes including Shift, Slide, Rise, Tilt, Roll, and HelT (helical twist); intra-base pair shapes including Shear, Stretch, Stagger, Buckle, ProT (propeller twist), and Opening; and minor groove shapes including MGW (minor groove width) and EP (electrostatic potential). The intra-base pair and minor groove features generate one output per sliding 5-mer, while the inter-base pair features generate two outputs per sliding 5-mer.

DNAshapeR also provides the estimates of four shape features (HelT, Roll, ProT, MGW) for methylated CpG dinucleotides. Since over 70% of all CpG sites are permanently methylated in the human genome (36), we have assumed all CpGs to be methylated and estimated their shape features as such. Thus, for 7-mers that include the “CG” dinucleotide, we would use the methylated shape data for HelT, Roll, ProT, and MGW, and use the normal shape data for all other features since DNAshapeR does not provide their estimates in the methylated state.

### Categorizing mutation rates into mutation classes

The *mutation class* of a given single nucleotide mutation represents the identities of its reference and alternative alleles. Following Aggarwala and Voight (3), we modeled each mutation class separately. Since we have folded complementary sequences, the central reference allele could only be A or C; thus, we have characterized a total of nine classes: A-to-C, A-to-G, A-to-T, C-to-A, C-to-G, C-to-T, CpG C-to-A, CpG C-to-G, CpG C-to-T. See Figure 1C (top) for an example of mutation class generation for a 5-mer mutation.

### Preparing model inputs from DNA sequence features

For the nucleotide sequence features, we have used a modified one-hot encoding scheme where we encode A as [0,0,0] and C, G, and T as [1,0,0], [0,1,0], and [0,0,1], respectively. We do not explicitly encode the mutated nucleotide (the middle position in the 7-mer), since this information does not change within a given mutation class. This results in 18 first-order features. Like Aggarwala and Voight, we allowed up to fourth-order interactions between nucleotide features. For the higher order interactions, we only considered nucleotides from different locations; same-location interactions would result in duplicated feature values. See Figure 1C (middle) for an example of how first- and second-order nucleotide features are calculated for a 5-mer mutation.

### Preparing model inputs from DNA shape features of a mutated nucleotide’s sequence context

We first included the measurements of all 14 DNA shape features available from DNAshapeR for all positions in the 7-mer context of a mutated nucleotide. This corresponds to 48 features for the reference 7-mer and the alternative 7-mer which we will refer to as ref and alt 7-mers, resulting in a total of 96 first-order features for our linear regression model. Then, we computed second-order features by 1) using the PolynomialFeatures function from *sklearn* with degree=2 and include_bias=False, resulting in a total of 4752 features (4158 after removing low variance features); alternatively, 2) we designed a custom function that computes polynomial terms but only included neighboring interactions, resulting in a total of 2282 features (2089 after removing low variance features). See Figure 1C (bottom) for more information.

Prior to making predictions, we have conducted min-max scaling of all shape features so that the maximum and minimum possible values for any first or second-order shape feature are (1, 0), while the geometric scale is preserved. This is done to 1) make linear regression models converge faster, and 2) ensure equal comparison of feature importance. Then, we removed features which had variances of less than 0.01 after min-max scaling as mentioned previously to ensure there are no low-variance features that may contribute minimally to model prediction.

Since adjacent sliding 5-mer windows may be intercorrelated, we have conducted a preliminary analysis of the shape features by calculating Pearson correlations between them (Figure S13). There is no observed strong correlation between the 14 different types of shapes, thus justifying our use of conventional regression model frameworks.

### Regression model building

Using the nucleotide sequence and shape features described above, we fit a separate regression model for each mutation class. For each model, we partitioned the data into training and test sets by computing mutation rates from all even-numbered and all odd-numbered chromosomes, and fit the model on the training data using 8-fold cross validation (CV); 8-fold is used as the total number of k-mers is divisible by 8 but not by 5 or 10. We then report the model’s performance on the separately held out test set. To eliminate redundant features, we used Lasso regularization, a technique that attempts to minimize the coefficients of redundant features to zero. Thus, given the feature matrix *X* (each row corresponds to a mutation *k*1 *→k*2 where *k*1 and *k*2 are the two 7-mers, and each column corresponds to a sequence or shape feature) and data vector *Y* (the mutation rates), we optimize the following objective function on the training data:

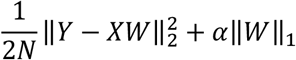

where *W* is the vector of model coefficients and α represents the Lasso regularization parameter: higher values of α implies stronger regularization, i.e., optimizing the objective function will require minimizing more coefficients toward zero. We searched for the optimal value of α from the following set of plausible values using 8-fold CV. We used a stepwise decreasing array of alpha values until decreasing the value of alpha increases the 8-fold CV validation MSE of our model; the set of values for α is determined by the following:

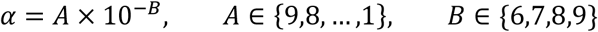

We used the “random” feature selection with a fixed seed to speed up our selections. We did not change the tolerance setting, but instead increased the maximum allowed iteration number so that the functions will eventually converge.

### Classification model building

Similar to the regression models, we first built the models using only the training data. We used decision tree classifiers, which iteratively identifies the most informative features for a given classification task. For each individual model, we used 8-fold CV as mentioned above to search for the optimal hyperparameter values from a set of hyperparameter combinations (minimum sample per split “min_samples_split”, minimum samples per leaf “min_samples_leaf”, the maximum number of features used “max_features”) using grid search (*sklearn*.GridSearchCV). These functions are available through *scikit-learn* (*sklearn*), a Python library of machine learning and statistical methods (40). We limited the depth of our decision trees to 2 to ensure easy interpretability. The best models were selected based on maximizing balanced accuracy, and model performances were evaluated based on ROC curves (false positive rate – true positive rate) and AUROC values based on predictions made on the held-out testing data. The optimization uses reductions in Gini impurity. For a binary classification, Gini impurity is calculated by:

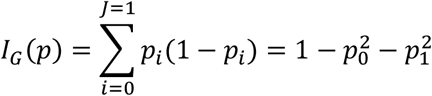

Intuitively, Gini impurity describes the imbalance in the distribution of labels in the subset. For binary classification with no class imbalance, a null model would produce a Gini impurity of 0.5, a 100% accurate model would produce a Gini impurity of 0, while a 0% accurate model would produce a Gini impurity of

### 1. Model interpretation

To compare the coefficient of each DNA shape feature across the regression models of different mutation classes, we first divided the value of each feature’s coefficient (*W*) by the range of values of the dependent variable (*Y*, the mutation rates) in that mutation class:

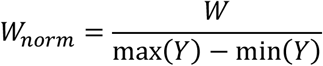

To compare the coefficient of each DNA shape feature across classification models, each feature is matched with a “feature importance” value, which is calculated by decreases in Gini impurities as mentioned above. For both situations, the scales of coefficient values are correlated with feature importance, thus we defined the absolute value of a feature coefficient as the *importance* of a feature.

For the regression models, we also considered how often a DNA shape feature was included in the optimized model. We computed two metrics. The first metric, *utilization*, is the count of a type of feature divided by the total number of features within the model. The second metric, *enrichment*, is the ratio of *utilization* in different importance bins of the model. By importance bins, we separated the regression model features into five or 10 equal-sized bins based on the percentile of feature importance of each feature; for example, the least important features would be assigned to bin 1, while the most important features would be assigned to bin 10. We classified features based on 1) which DNA shape they belong to, 2) their relative location in the 7-mer, and 3) their location on the reference or the alternative 7-mer. We also included the exact model descriptions of the classification models including the identities of the nodes as well as decision boundaries (Figure 4D, Supplemental figure 13).

For orthogonal validation of our classifications models, individual 7-mers were hand-curated by comparing their shape values to the decision boundaries defined by our decision tree models.

### Transcription factor analyses

Transcription factor binding data was retrieved from Kheradpour and Kellis, 2014 (31). The downloaded data were processed on a per-motif basis. For each motif, we processed the data to include 1) DNA shape values across all hits for a particular motif, with 2 bp of extended sequence upstream and downstream to avoid the lack of edge prediction in DNAshapeR, and 2) phase 3 1KG variants that fall within these binding motifs, their relative distance from the first DNA base (positive strand) of the corresponding motif, as well as their local 5-mer sequence context for DNA shape estimation. Transcription factor family classification data was retrieved from Wingender et al. (41), and the processed data was provided in Table S15.

For our TFBS analyses, we conducted all of our analyses on a per-motif and per-position basis. We first calculated DNA shape values from the mutated k-mers. Then, for each position of a given motif, we calculated 1) Kolmogorov-Smirnov test statistic between the shape distribution of the motif hits and the shape distribution of the mutated 5-mers, 2) position-specific observed mutation rate by dividing the number of observed mutations that fall on this position of the motif by the total number of motif hits in the genome, and 3) Shannon entropy by calling the scipy.stats.entropy function on the position frequency matrix of a motif with a default probability distribution of [0.25, 0.25, 0.25, 0.25] mapping to A/C/G/T. We then calculated Spearman rank correlations between these three metrics measured on different positions of each motif, on a per-motif basis. For our final analyses, we averaged correlation values across all unique motifs for each unique TF so that analyses were done on a per-TF instead of a per-motif basis.

## Model implementation

We have used Python and Jupyterlab for coding tasks; the library *scikit-learn* (*sklearn*) (36) and *scipy* were used for our model building exercises, with certain tasks run on an Ubuntu terminal. The R package DNAshapeR was used to obtain DNA shape features.

## Supporting information

Oversized supplemental tables

## Description of Supplemental Data

The supplemental data contains 13 figures and 15 tables.

## Acknowledgements

We thank Benjamin F. Voight, Ph.D. for kindly providing the input data and guidance on data processing for this manuscript. We thank the Samee Lab members for sharing their comments during the development of this project.

## Web Resources

The R package DNAshapeR can be accessed at http://tsupeichiu.github.io/DNAshapeR/; the R script for extracting reference 7-mer DNA shape spreadsheets from DNAshapeR can be accessed at https://github.com/ZnL-BCM/DNAshapeR_reference. Data from the 1000 Genomes Project can be accessed at https://www.internationalgenome.org/home.

## Data and Code Availability

We obtained the mutation rate data by requesting Benjamin F. Voight, Ph.D., the corresponding author of the Aggarwala and Voight publication (3). All other data, as well as the example code, notebooks, and certain results used for this publication, are available in a repository on the Samee Lab GitHub at https://github.com/sameelab/mutprediction-with-shape.

