## Supplemental information for "Mutation Rate Variations in the Human Genome are Encoded in DNA Shape"

**Supplementary Figures**

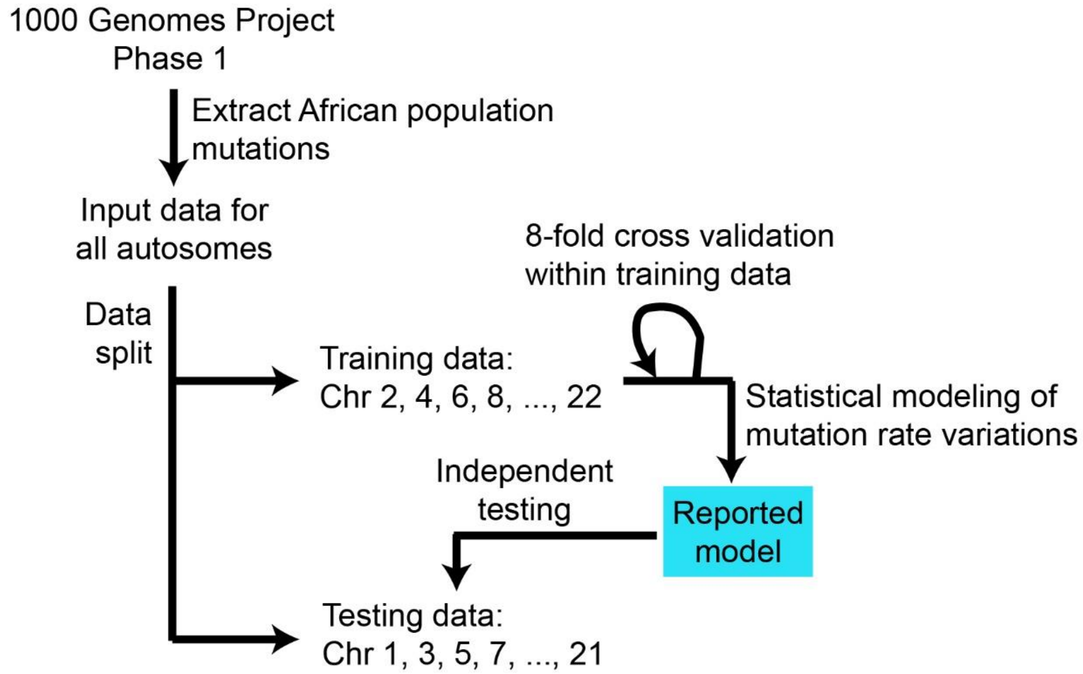


**Figure S1**. Flowchart of data usage and modeling strategies for our models.

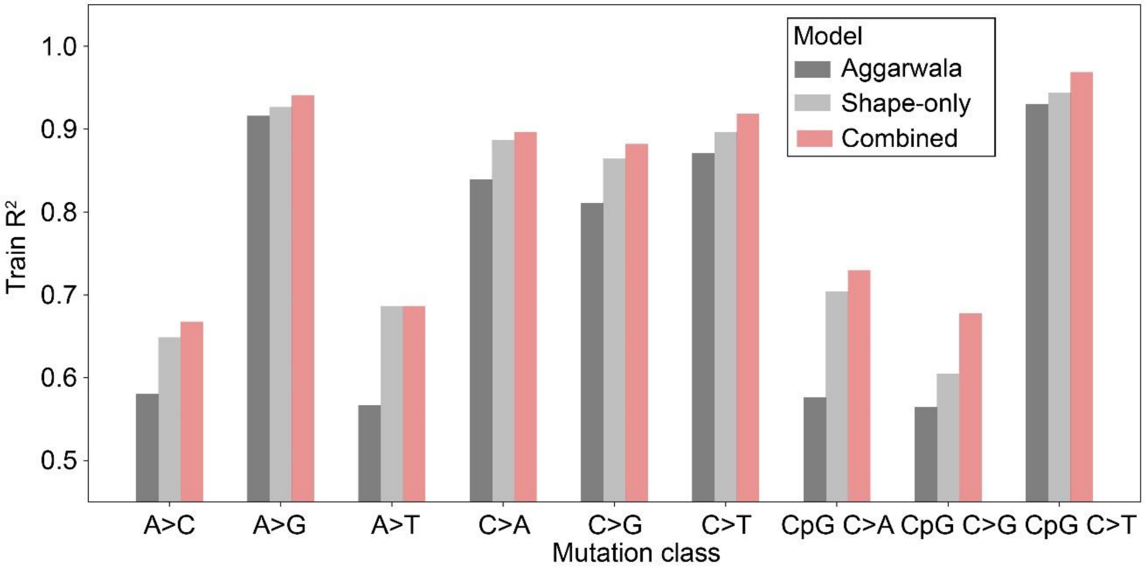


**Figure S2**. Bar chart showing training $R^2$ values of our best performing model (red), the Aggarwala and Voight model (black), and the Lasso-based feature selection fourth order nucleotide interaction model (grey).

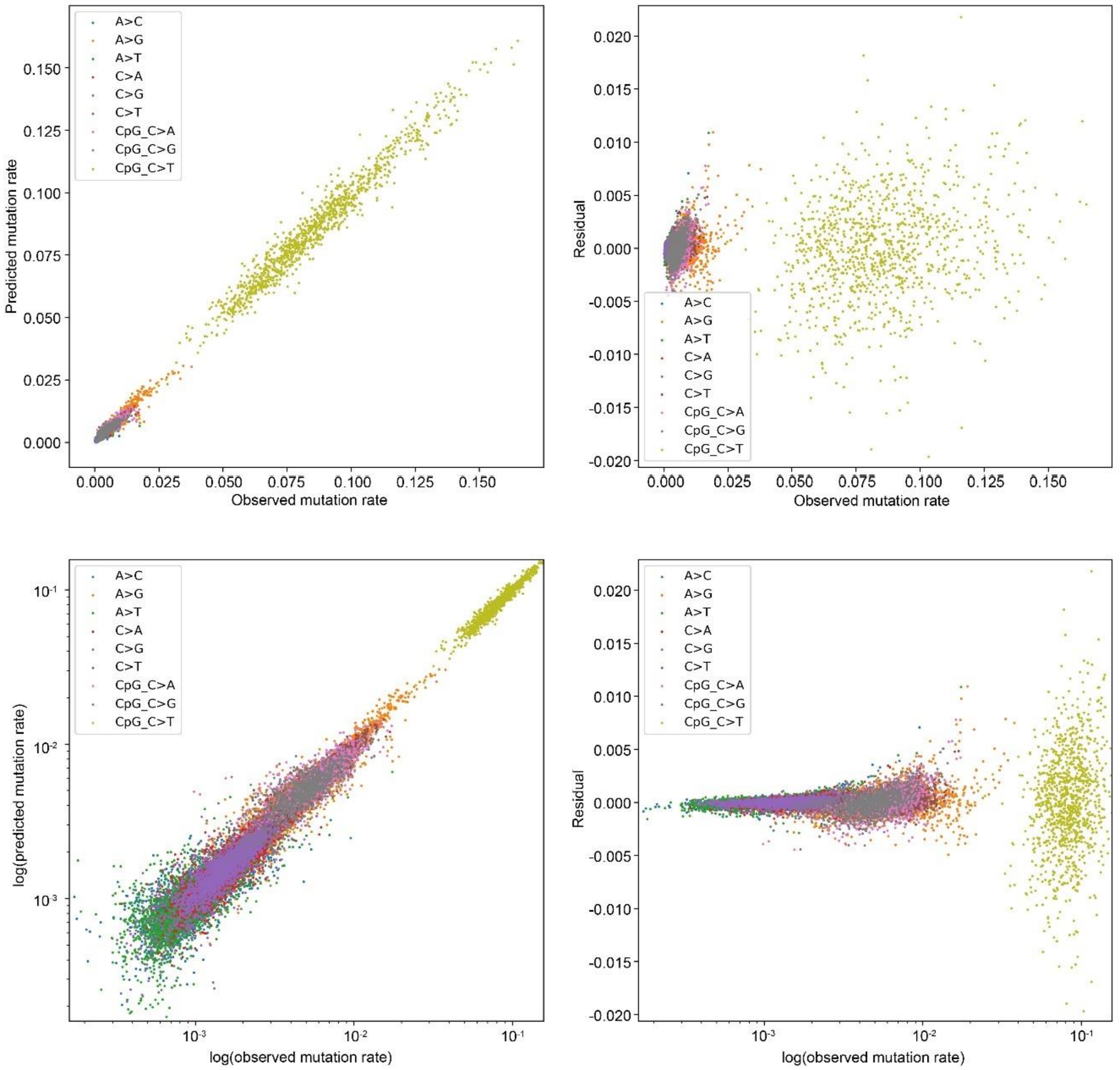


**Figure S3**. Scatterplots (left) and residual plots (right) of predictions from our best performing model on the training data, in linear scale (top) and logarithmic scale (bottom). All x-axes are observed mutation rates, while y-axes are model-predicted mutation rates (left) and residuals (right).

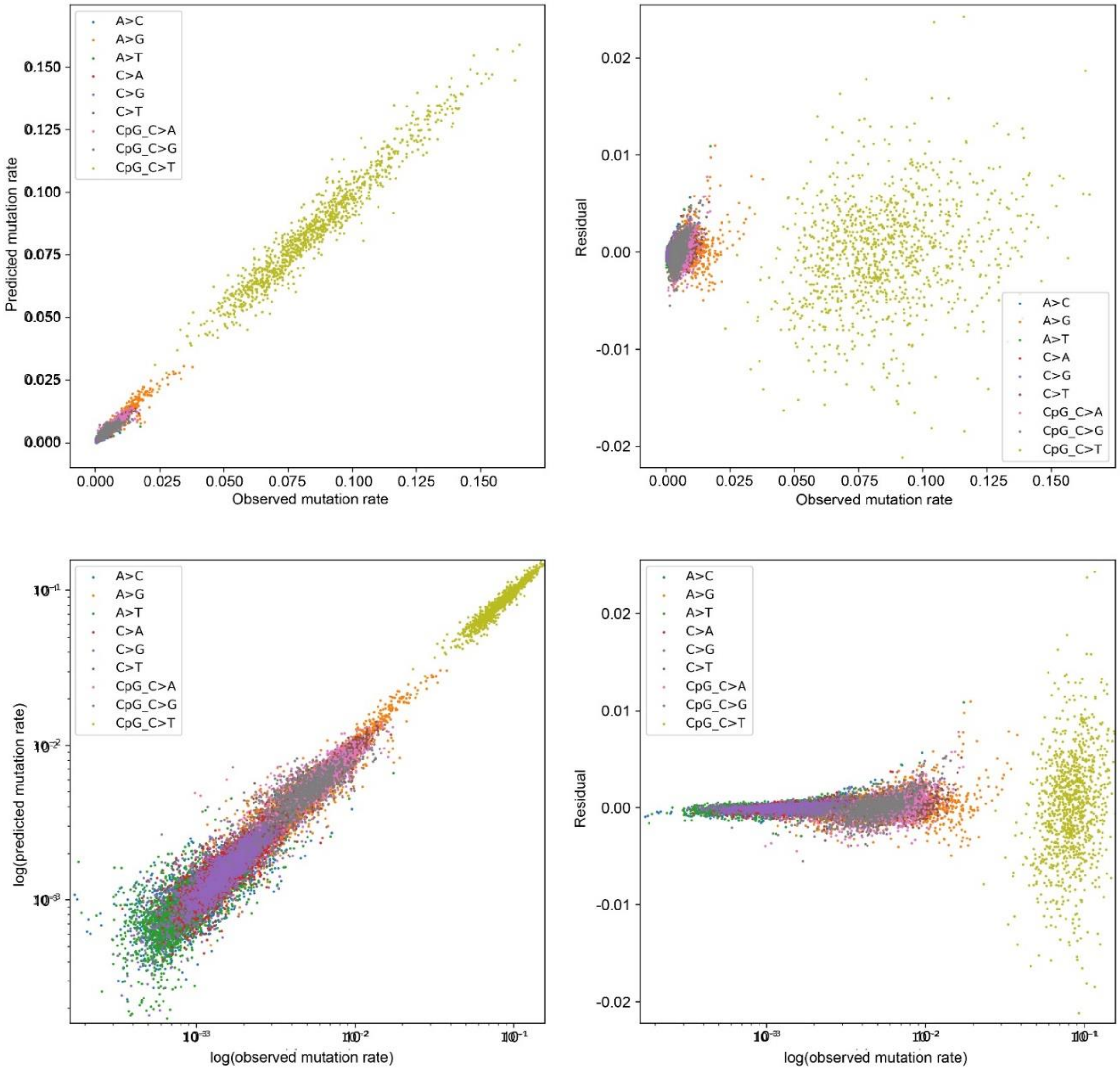


**Figure S4**. Scatterplots (left) and residual plots (right) of predictions from our second-order shape model on the training data, in linear scale (top) and logarithmic scale (bottom). All x-axes are observed mutation rates, while y-axes are model-predicted mutation rates (left) and residuals (right).

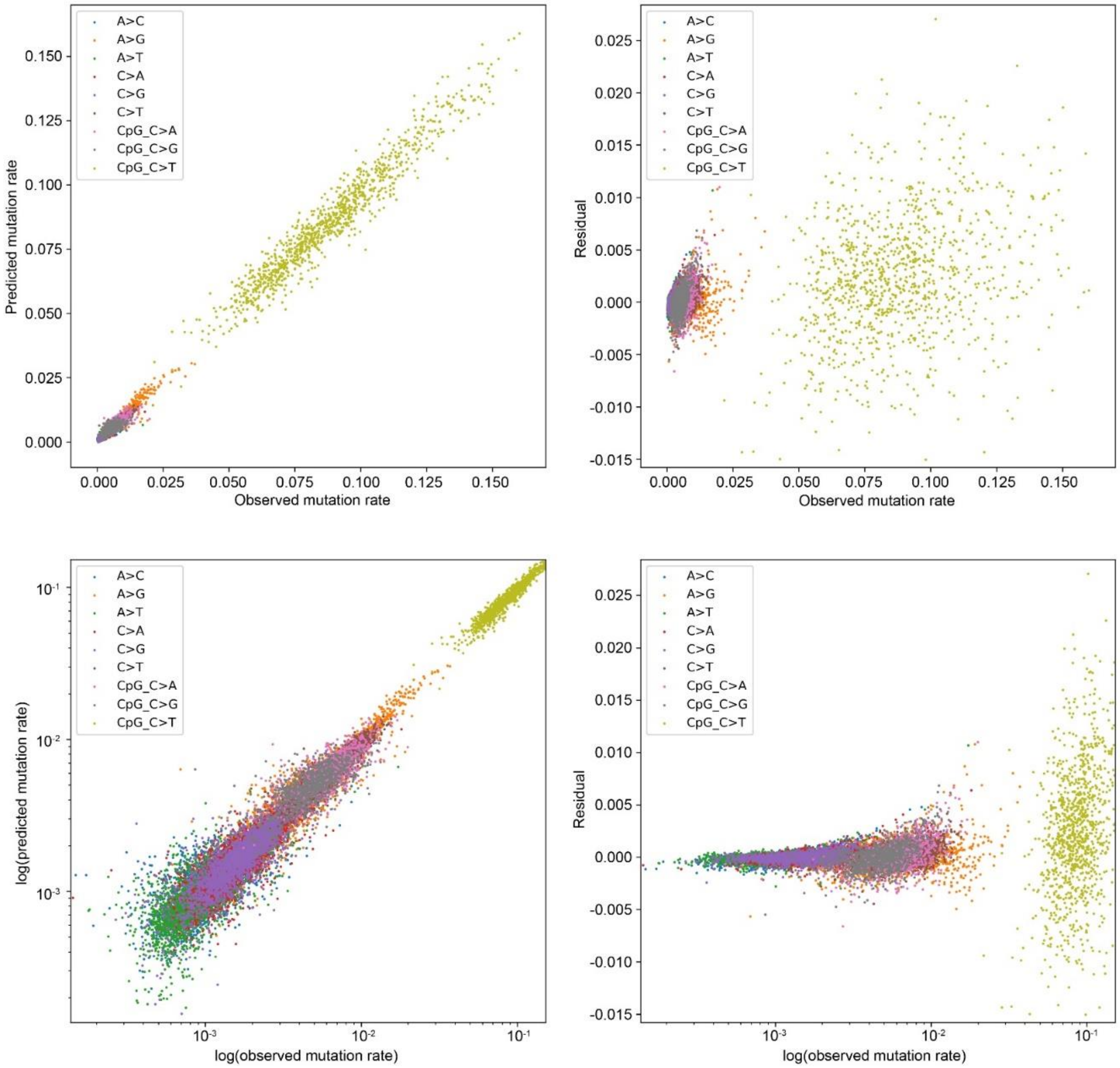


**Figure S5**. Scatterplots (left) and residual plots (right) of predictions from our second-order shape model on the testing data, in linear scale (top) and logarithmic scale (bottom). All x-axes are observed mutation rates, while y-axes are model-predicted mutation rates (left) and residuals (right).

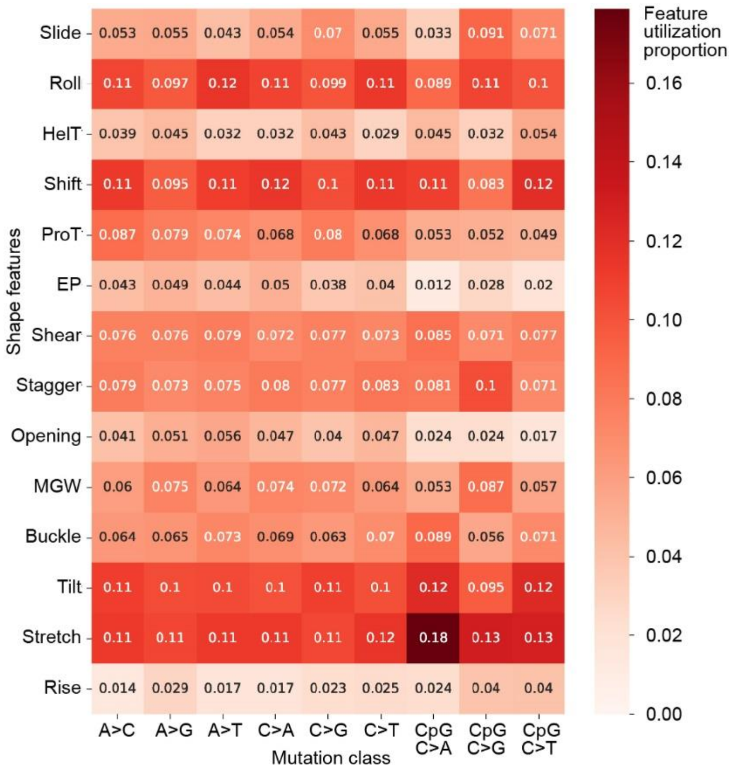


**Figure S6.** Relative utilization of 14 shape features across regression models in all nine mutation classes using all features.

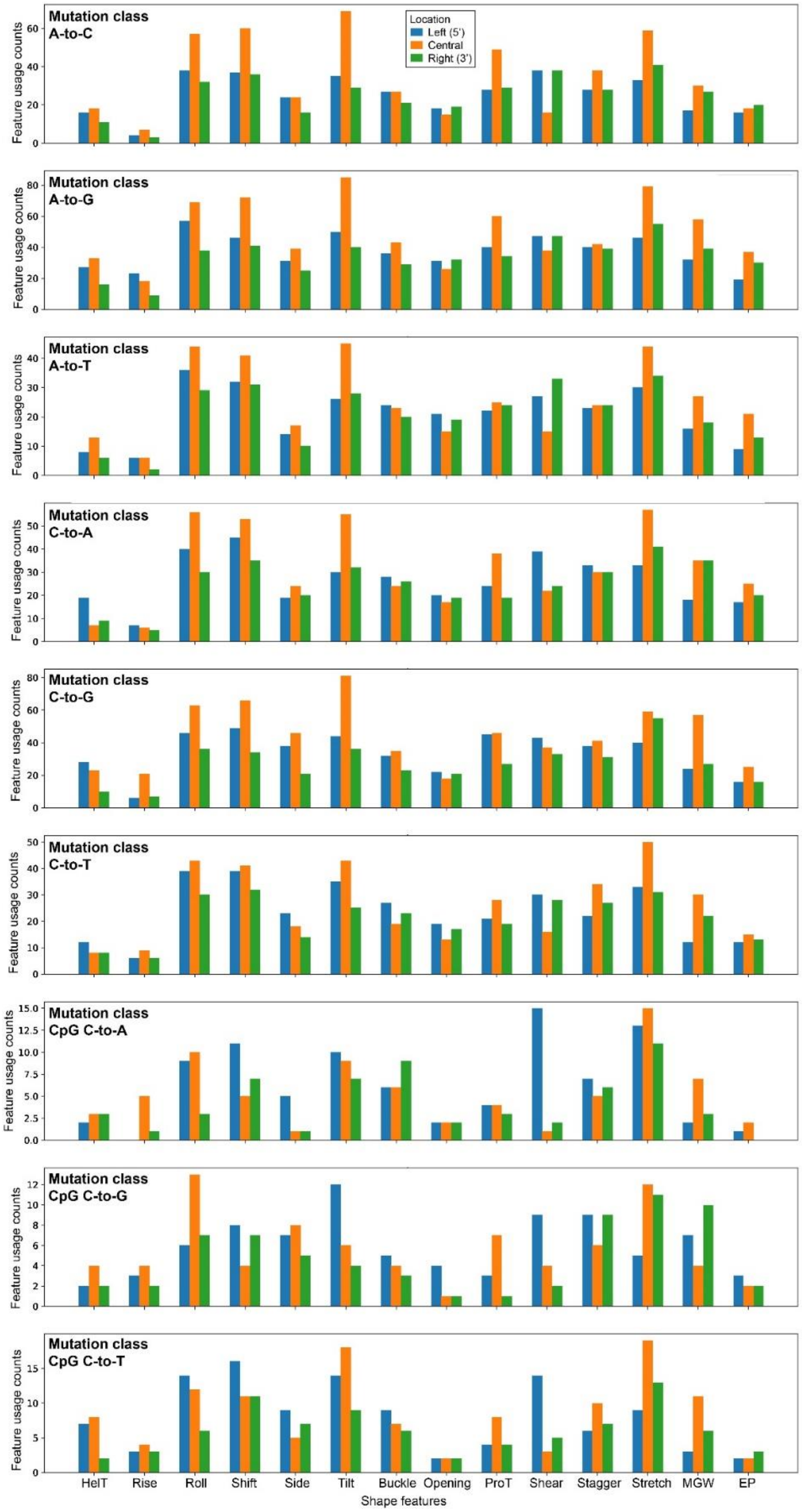


**Figure S7**. Bar charts of feature utilization based on interactions of shape features with particular locations for all nine mutation classes. The x-axes denote mutation classes and is shared across all nine subplots, and the y-axes denote feature usage in a particular model. The figure legend is shared across all nine subplots.

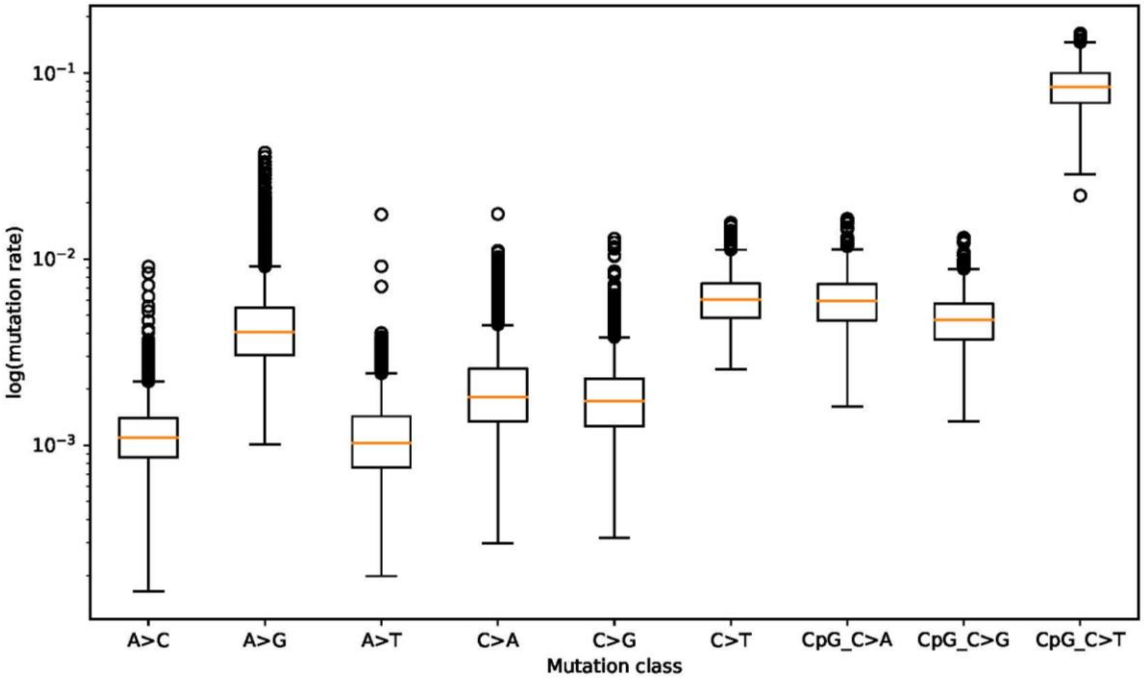


**Figure S8**. Boxplot of mutation rate variations for each of the nine mutation classes.

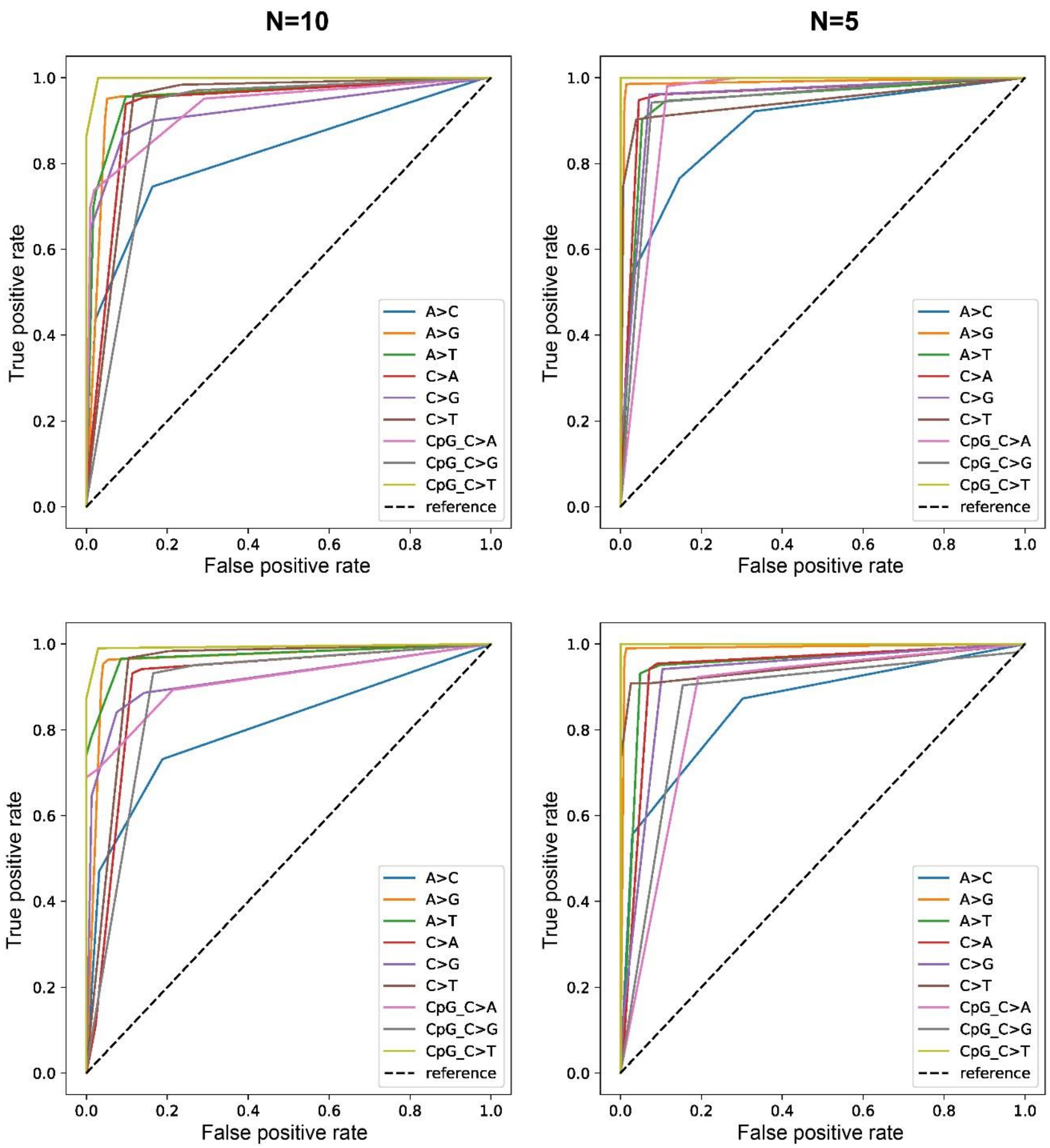


**Figure S9**. Receiver operating curves (ROC) for decision tree modeling of high versus low mutation rates with N in {5, 10} in all nine mutation classes. The subplots represent ROC curves on training (top) and testing (bottom) data, as well as values of N equal to 10 (left) or 5 (right). The black dashed lines are diagonal reference lines.

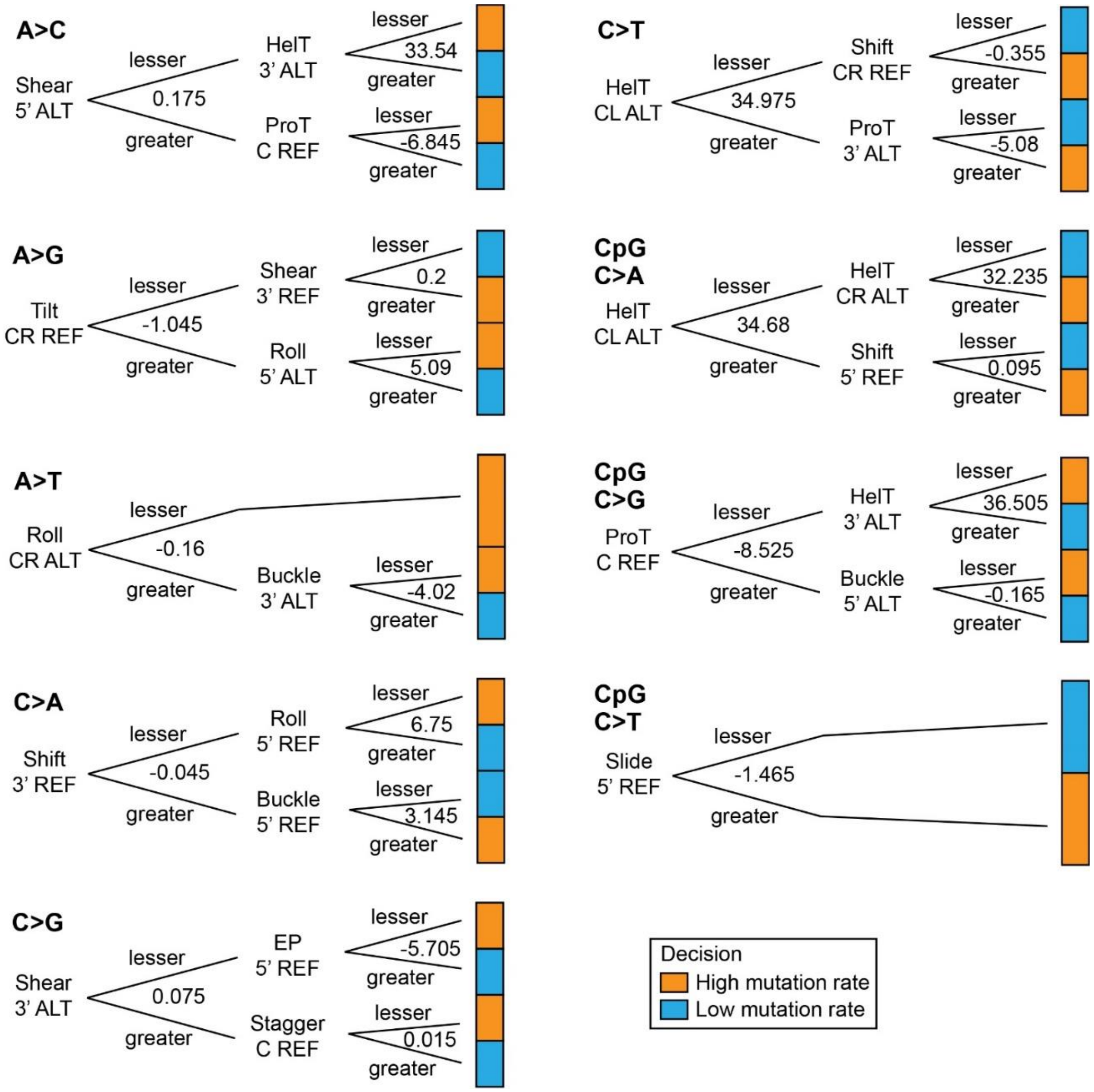


**Figure S10**. Illustrations of decision tree architectures with N=10 for the mutation classes A) A-to-T, B) C-to-A, C) C-to-G, D) C-to-T, E) CpG C-to-A, F) CpG C-to-G, G) CpG C-to-T.

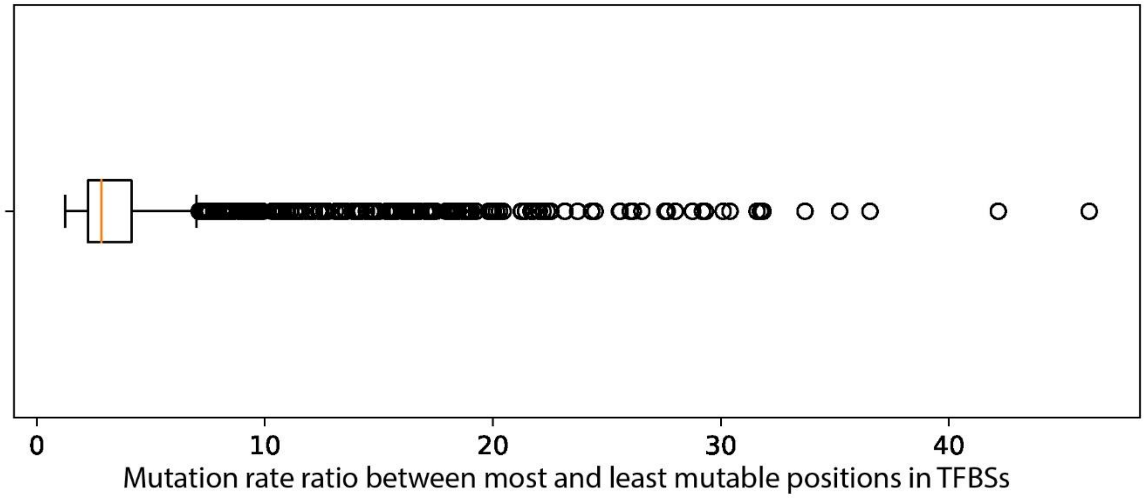


**Figure S11**. Ratio between the mutation rates of the most and least mutable positions in TFBSs which harbor single nucleotide mutations.

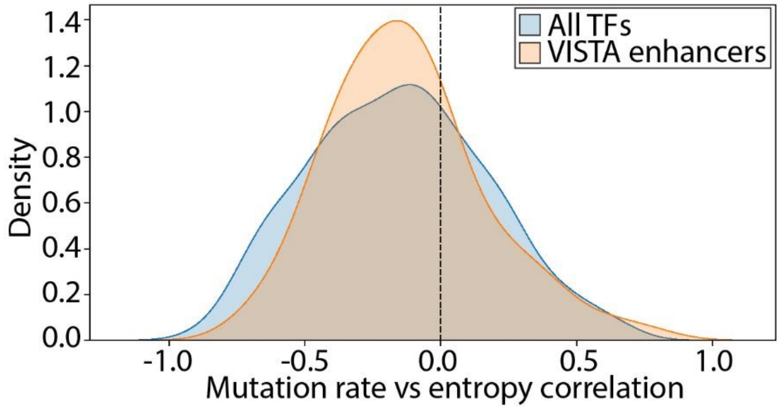

**Figure S12**. Distribution of spearman rank correlation values between per-position entropy values and KS statistics of helical twist shape changes for all transcription factors and transcription factors that commonly occur in VISTA enhancer regions.

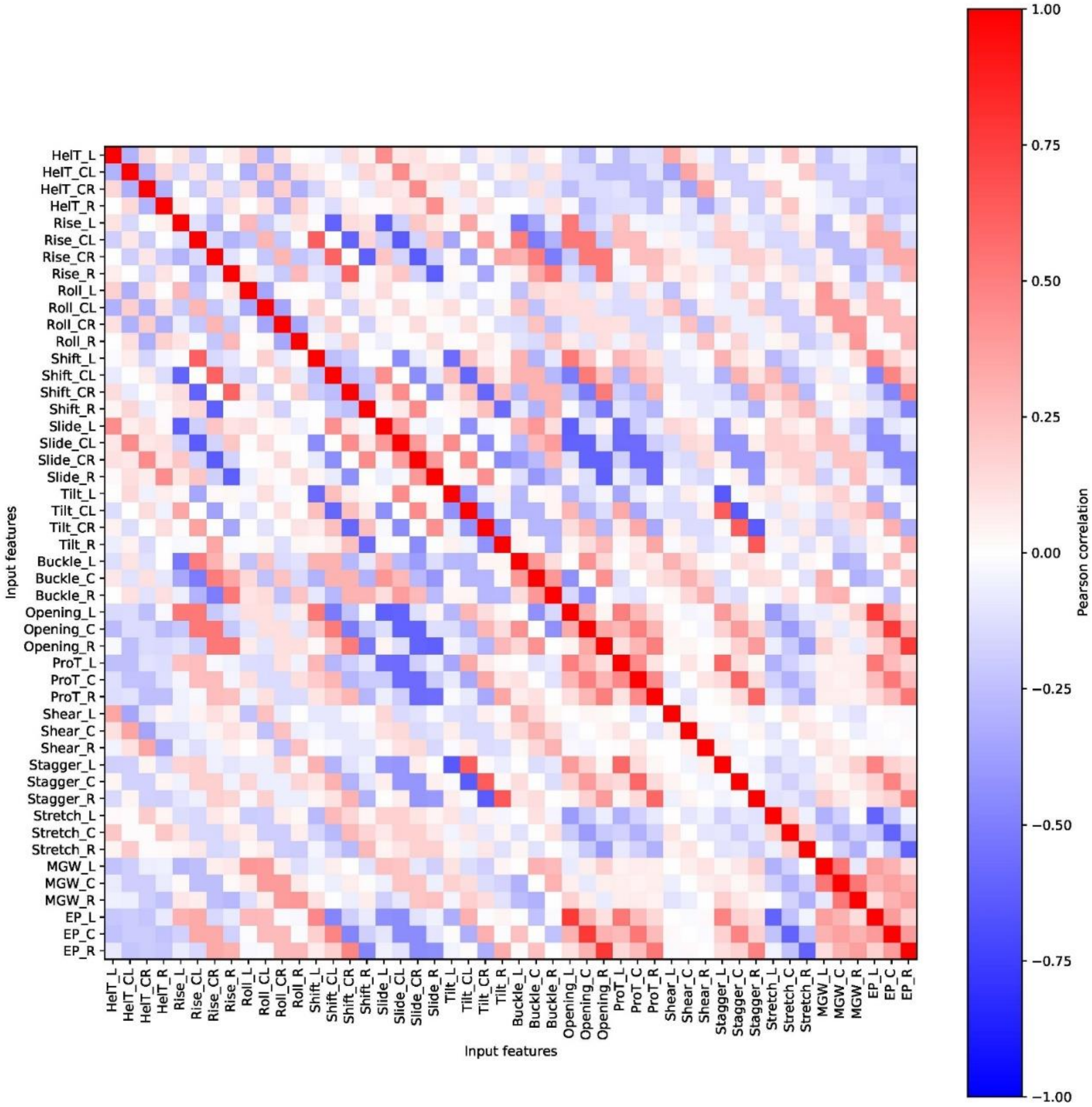


**Figure S13**. Heatmap of Pearson correlations of all 7-mer input DNA shape features.

### Supplemental Tables

#### Table legends

**Tables S3-4** record model parameters for two of our regression models that are worth mentioning. The horizontal axes represent the nine mutation classes, while the vertical axes represent feature identities. Each first-order shape feature is written as ShapeIdentity_location[_r], while an asterisk (*) signals that the feature is an interaction of two shape features. The possible locations include 5'/left (L), central left (CL), central (C), central right (CR), and 3'/right (R), see Methods section for more information. If "_r" is present, then this shape is on the alternative 7-mer. Each nucleotide feature is written exactly as the nucleotide identities it represents; note that there are only six nucleotides as the central nucleotide is not encoded (see Methods).

**Table S5** record Benjamin-Hochberg false discovery rate (FDR-BH)-adjusted p-values of the binomial tests. When conducting binomial tests, the features were classified into 10 importance bins based on the ranking of their feature importance (see Methods).

**Tables S11-12** record Spearman rank correlation values of KS statistics of DNA shape changes versus observed mutation rates across S11) all TFBS motifs, or S12) motifs that commonly occur in VISTA enhancer regions as defined by Table S10. Note that correlation values from motifs of the same transcription factors were averaged for the main figures.

**Table S15** record the corresponding table of transcription factor binding site motifs, transcription factors, transcription factor family definitions, as well as whether each motif is common within VISTA enhancer regions.

Please note that **Tables S3-5**, **S11-12**, and **S15** are oversized and will be included in a separate xlsx document.

**Table S1**. Model performance across various combinations of nucleotide and shape features on training data. The “maximum achievable $R^2$” column were calculated by directly comparing test data with itself and is thus always 1. The “Aggarwala” columns describe performances of the current state of the art model. The remaining columns represent models that we fitted. Regarding abbreviations, "sh" and "sc" represent shape and nucleotide (sequence context) features, and the "neibr" suffix represents that only neighboring interactions were included (see Methods). The mean and median values were computed based on $R^2$ values of the nine individual models and were not computed for performance changes to avoid ambiguity.

| **Mutation class** | **Maximum achievable $R^2$** | **Aggarwala** | **sh1** | **sh2neibr** | **sc4** | **sh1_sc1** | **sh1_sc2** | **sh1_sc3** | **sh1_sc4** | **sh2neibr_sc1** |
|---|---|---|---|---|---|---|---|---|---|---|
| A>C | 1.000 | 0.580 | 0.310 | 0.648 | 0.653 | 0.355 | 0.476 | 0.572 | 0.677 | 0.649 |
| A>G | 1.000 | 0.916 | 0.608 | 0.926 | 0.938 | 0.650 | 0.790 | 0.892 | 0.940 | 0.924 |
| A>T | 1.000 | 0.567 | 0.457 | 0.686 | 0.606 | 0.482 | 0.536 | 0.597 | 0.639 | 0.635 |
| C>A | 1.000 | 0.840 | 0.580 | 0.886 | 0.908 | 0.626 | 0.733 | 0.858 | 0.904 | 0.896 |
| C>G | 1.000 | 0.811 | 0.435 | 0.864 | 0.867 | 0.496 | 0.619 | 0.757 | 0.869 | 0.882 |
| C>T | 1.000 | 0.871 | 0.716 | 0.896 | 0.915 | 0.738 | 0.825 | 0.887 | 0.916 | 0.908 |
| CpG_C>A | 1.000 | 0.576 | 0.592 | 0.704 | 0.753 | 0.629 | 0.624 | 0.745 | 0.755 | 0.699 |
| CpG_C>G | 1.000 | 0.565 | 0.491 | 0.604 | 0.690 | 0.499 | 0.531 | 0.687 | 0.696 | 0.557 |
| CpG_C>T | 1.000 | 0.930 | 0.878 | 0.943 | 0.964 | 0.901 | 0.932 | 0.956 | 0.966 | 0.946 |
| **Mean** | 1.000 | 0.739 | 0.563 | 0.796 | 0.810 | 0.597 | 0.674 | 0.772 | 0.818 | 0.788 |
| **Median** | 1.000 | 0.811 | 0.580 | 0.864 | 0.867 | 0.626 | 0.624 | 0.757 | 0.869 | 0.882 |

| **Mutation class** | **sh2neibr_sc2** | **sh2neibr_sc3** | **sh2neibr_sc4** |
|---|---|---|---|
| A>C | 0.628 | 0.586 | 0.668 |
| A>G | 0.918 | 0.927 | 0.941 |
| A>T | 0.635 | 0.605 | 0.658 |
| C>A | 0.894 | 0.893 | 0.908 |
| C>G | 0.844 | 0.770 | 0.872 |
| C>T | 0.901 | 0.893 | 0.919 |
| CpG_C>A | 0.663 | 0.730 | 0.737 |
| CpG_C>G | 0.531 | 0.660 | 0.676 |
| CpG_C>T | 0.944 | 0.957 | 0.969 |
| **Mean** | 0.773 | 0.780 | 0.816 |
| **Median** | 0.844 | 0.770 | 0.872 |

**Table S2**. Model performance across various combinations of nucleotide and shape features on test data. The "maximum achievable $R^2$" column were calculated by directly comparing test data with the training data. The "Aggarwala" columns describe performances of the current state of the art model. The remaining columns represent models that we fitted. Regarding abbreviations, "sh" and "sc" represent shape and nucleotide (sequence context) features, and the "neibr" suffix represents that only neighboring interactions were included (see Methods). The mean and median values were computed based on $R^2$ values of the nine individual models and were not computed for performance changes to avoid ambiguity.

| Mutation class | Maximum achievable $R^2$ | Aggarwala | sh1 | sh2neibr | sc4 | sh1_sc1 | sh1_sc2 | sh1_sc3 | sh1_sc4 | sh2neibr_sc1 |
|---|---|---|---|---|---|---|---|---|---|---|
| A>C | 0.666 | 0.580 | 0.300 | 0.579 | 0.582 | 0.341 | 0.455 | 0.533 | 0.586 | 0.579 |
| A>G | 0.961 | 0.920 | 0.615 | 0.922 | 0.929 | 0.659 | 0.800 | 0.893 | 0.930 | 0.921 |
| A>T | 0.750 | 0.605 | 0.489 | 0.673 | 0.591 | 0.514 | 0.569 | 0.603 | 0.610 | 0.650 |
| C>A | 0.905 | 0.840 | 0.579 | 0.868 | 0.877 | 0.625 | 0.731 | 0.845 | 0.878 | 0.873 |
| C>G | 0.867 | 0.819 | 0.451 | 0.834 | 0.841 | 0.513 | 0.628 | 0.758 | 0.842 | 0.844 |
| C>T | 0.864 | 0.875 | 0.714 | 0.870 | 0.878 | 0.736 | 0.817 | 0.871 | 0.878 | 0.873 |
| CpG_C>A | 0.534 | 0.553 | 0.559 | 0.635 | 0.610 | 0.579 | 0.574 | 0.618 | 0.609 | 0.626 |
| CpG_C>G | 0.417 | 0.531 | 0.440 | 0.517 | 0.527 | 0.449 | 0.491 | 0.529 | 0.526 | 0.480 |
| CpG_C>T | 0.934 | 0.932 | 0.867 | 0.925 | 0.931 | 0.889 | 0.916 | 0.930 | 0.932 | 0.926 |
| **Mean** | 0.766 | 0.740 | 0.557 | 0.758 | 0.752 | 0.589 | 0.665 | 0.731 | 0.755 | 0.752 |
| **Median** | 0.864 | 0.819 | 0.559 | 0.834 | 0.841 | 0.579 | 0.628 | 0.758 | 0.842 | 0.844 |

| Mutation class | sh2neibr_sc2 | sh2neibr_sc3 | sh2neibr_sc4 |
|---|---|---|---|
| A>C | 0.569 | 0.543 | 0.592 |
| A>G | 0.916 | 0.924 | 0.931 |
| A>T | 0.653 | 0.616 | 0.631 |
| C>A | 0.872 | 0.875 | 0.882 |
| C>G | 0.824 | 0.768 | 0.844 |
| C>T | 0.874 | 0.876 | 0.882 |
| CpG_C>A | 0.603 | 0.615 | 0.611 |
| CpG_C>G | 0.493 | 0.530 | 0.524 |
| CpG_C>T | 0.927 | 0.930 | 0.934 |
| **Mean** | 0.748 | 0.742 | 0.759 |
| **Median** | 0.824 | 0.768 | 0.844 |

**Table S6**. Area under receiver operating curve (AUROC) values for decision tree models with N in {5, 10} and all mutation classes on training and test data. The horizontal axis denotes the values of N, which corresponds to the proportion of k-mers with the highest and lowest N percent of mutation rates subsequently used for the classification task.

| **Mutation class** | **10%_train** | **10%_test** | **5%_train** | **5%_test** |
|---|---|---|---|---|
| A>C | 0.821 | 0.804 | 0.883 | 0.857 |
| A>G | 0.953 | 0.962 | 0.988 | 0.990 |
| A>T | 0.956 | 0.972 | 0.943 | 0.948 |
| C>A | 0.928 | 0.908 | 0.957 | 0.939 |
| C>G | 0.923 | 0.917 | 0.950 | 0.923 |
| C>T | 0.931 | 0.938 | 0.944 | 0.949 |
| CpG_C>A | 0.932 | 0.913 | 0.940 | 0.866 |
| CpG_C>G | 0.895 | 0.889 | 0.935 | 0.868 |
| CpG_C>T | 0.998 | 0.993 | 1.000 | 1.000 |

**Table S7**. Confusion matrices of decision tree models on training data.

| | | N=10 Predicted low rate | N=10 Predicted high rate | N=5 Predicted low rate | N=5 Predicted high rate |
|---|---|---|---|---|---|
| **A>C** | Actual low rate | 343 | 67 | 175 | 30 |
| | Actual high rate | 104 | 306 | 48 | 157 |
| **A>G** | Actual low rate | 389 | 21 | 202 | 3 |
| | Actual high rate | 20 | 390 | 3 | 202 |
| **A>T** | Actual low rate | 370 | 40 | 194 | 11 |
| | Actual high rate | 18 | 392 | 20 | 185 |
| **C>A** | Actual low rate | 278 | 30 | 147 | 7 |
| | Actual high rate | 19 | 289 | 8 | 146 |
| **C>G** | Actual low rate | 280 | 28 | 143 | 11 |
| | Actual high rate | 41 | 267 | 6 | 148 |
| **C>T** | Actual low rate | 272 | 36 | 148 | 6 |
| | Actual high rate | 12 | 296 | 15 | 139 |
| **CpG C>A** | Actual low rate | 101 | 2 | 46 | 6 |
| | Actual high rate | 27 | 76 | 1 | 51 |
| **CpG C>G** | Actual low rate | 85 | 18 | 48 | 4 |
| | Actual high rate | 5 | 98 | 3 | 49 |
| **CpG C>T** | Actual low rate | 100 | 3 | 52 | 0 |
| | Actual high rate | 0 | 103 | 0 | 52 |

**Table S8**. Confusion matrices of decision tree models on test data.

| | | N=10 Predicted low rate | N=10 Predicted high rate | N=5 Predicted low rate | N=5 Predicted high rate |
|---|---|---|---|---|---|
| **A>C** | Actual low rate | 333 | 77 | 176 | 29 |
| | Actual high rate | 110 | 300 | 64 | 141 |
| **A>G** | Actual low rate | 393 | 17 | 202 | 3 |
| | Actual high rate | 19 | 391 | 2 | 203 |
| **A>T** | Actual low rate | 375 | 35 | 195 | 10 |
| | Actual high rate | 14 | 396 | 14 | 191 |
| **C>A** | Actual low rate | 273 | 35 | 143 | 11 |
| | Actual high rate | 21 | 287 | 9 | 145 |
| **C>G** | Actual low rate | 285 | 23 | 138 | 16 |
| | Actual high rate | 49 | 259 | 9 | 145 |
| **C>T** | Actual low rate | 276 | 32 | 150 | 4 |
| | Actual high rate | 10 | 298 | 14 | 140 |
| **CpG C>A** | Actual low rate | 100 | 3 | 42 | 10 |
| | Actual high rate | 30 | 73 | 4 | 48 |
| **CpG C>G** | Actual low rate | 86 | 17 | 44 | 8 |
| | Actual high rate | 7 | 96 | 5 | 47 |
| **CpG C>T** | Actual low rate | 100 | 3 | 52 | 0 |
| | Actual high rate | 1 | 102 | 0 | 52 |

**Table S9**. Summary of decision tree interpretation results of all 7-mers under certain motifs as described in Aggarwala and Voight (3). The tree set used was for N=5. Agreement denotes the percentage of 7-mers in each sequence motif where our decision trees reached consensus with the expected effects on mutation probability for that particular motif.

| sequence motif | mutation class | Total # of 7-mers | expected effect on mutation probability | Agreement between Aggarwala and our model |
|---|---|---|---|---|
| N[A/C/G][C/G/T]CGCG | CpG C-to-T | 36 | lower | 100% |
| NTACG[C/G][A/C/G] | CpG C-to-T | 24 | higher | 83.30% |
| poly-A and poly-T combinations | A-to-T | 2 | higher | 50% |
| A4 | A-to-G | 208 | lower | 73.10% |
| [C/T]CAAT[C/G/T]N | A-to-G | 24 | higher | 100% |

**Table S10**. Decision tree interpretations of six 7-mers under certain motifs as described in Aggarwala and Voight (3). The tree set used was for N=5. Our decision trees reached consensus with the expected effects on mutation probability in all six instances.

| 7-mer | mutation class | sequence motif | expected effect on mutation probability | Root feature | Branch feature |
|---|---|---|---|---|---|
| CGGCGCG | CpG C-to-T | N[A/C/G][C/G/T]CGCG | lower | Slide_5'_ref | NaN |
| ATACGGG | CpG C-to-T | NTACG[C/G][A/C/G] | higher | Slide_5'_ref | Roll_3'_ref |
| AAAATTT | A-to-T | poly-A and poly-T combinations | higher | Roll_CR_alt | Roll_3'_ref |
| AAAAGAG | A-to-G | A4 | lower | Tilt_CR_ref | Shear_C_alt |
| GTTAAAA | A-to-G | A4 | lower | Tilt_CR_ref | Opening_L_alt |
| CCAATGG | A-to-G | [C/T]CAAT[C/G/T]N | higher | Tilt_CR_ref | Opening_5'_alt |

**Table S13**. Model performance between models that include or exclude non-neighboring shape interactions on training data. The "maximum achievable $R^2$" column were calculated by directly comparing test data with itself and is always 1. The "Aggarwala" columns describe performances of the current state of the art model. The remaining columns represent models that we fitted. Regarding abbreviations, "sh" and "sc" represent shape and nucleotide (sequence context) features, and the "neibr" suffix represents that only neighboring interactions were included (see Methods). The mean and median values were computed based on $R^2$ values of the nine individual models and were not computed for performance changes to avoid ambiguity.

| Mutation class | Maximum achievable $R^2$ | sh1 | sh2 | sh2neibr | sc2 | sc2neibr | sc3 | sc3neibr | sc4 | sc4neibr | sh1_sc2 | sh1_sc2neibr |
|---|---|---|---|---|---|---|---|---|---|---|---|---|
| A>C | 1.000 | 0.310 | 0.598 | 0.648 | 0.427 | 0.335 | 0.569 | 0.427 | 0.653 | 0.483 | 0.476 | 0.407 |
| A>G | 1.000 | 0.608 | 0.923 | 0.926 | 0.746 | 0.641 | 0.894 | 0.729 | 0.938 | 0.744 | 0.790 | 0.712 |
| A>T | 1.000 | 0.457 | 0.670 | 0.686 | 0.510 | 0.459 | 0.592 | 0.521 | 0.606 | 0.563 | 0.536 | 0.526 |
| C>A | 1.000 | 0.580 | 0.911 | 0.886 | 0.661 | 0.598 | 0.825 | 0.696 | 0.908 | 0.761 | 0.733 | 0.681 |
| C>G | 1.000 | 0.435 | 0.868 | 0.864 | 0.558 | 0.458 | 0.755 | 0.605 | 0.867 | 0.711 | 0.619 | 0.556 |
| C>T | 1.000 | 0.716 | 0.898 | 0.896 | 0.792 | 0.716 | 0.885 | 0.806 | 0.915 | 0.827 | 0.825 | 0.716 |
| CpG_C>A | 1.000 | 0.592 | 0.695 | 0.704 | 0.632 | 0.557 | 0.738 | 0.616 | 0.753 | 0.657 | 0.624 | 0.568 |
| CpG_C>G | 1.000 | 0.491 | 0.620 | 0.604 | 0.522 | 0.425 | 0.656 | 0.497 | 0.690 | 0.549 | 0.531 | 0.509 |
| CpG_C>T | 1.000 | 0.878 | 0.935 | 0.943 | 0.922 | 0.862 | 0.953 | 0.900 | 0.964 | 0.924 | 0.932 | 0.922 |
| **Mean** | 1.000 | 0.563 | 0.791 | 0.796 | 0.641 | 0.561 | 0.763 | 0.644 | 0.810 | 0.691 | 0.674 | 0.622 |
| **Median** | 1.000 | 0.580 | 0.868 | 0.864 | 0.632 | 0.557 | 0.755 | 0.616 | 0.867 | 0.711 | 0.624 | 0.568 |

| Mutation class | sh1_sc3 | sh1_sc3neibr | sh1_sc4 | sh1_sc4neibr | sh2neibr_sc2 | sh2neibr_sc2neibr | sh2neibr_sc3 | sh2neibr_sc3neibr | sh2neibr_sc4 | sh2neibr_sc4neibr |
|---|---|---|---|---|---|---|---|---|---|---|
| A>C | 0.572 | 0.473 | 0.677 | 0.547 | 0.628 | 0.643 | 0.586 | 0.670 | 0.668 | 0.612 |
| A>G | 0.892 | 0.796 | 0.940 | 0.853 | 0.918 | 0.916 | 0.927 | 0.913 | 0.941 | 0.914 |
| A>T | 0.597 | 0.564 | 0.639 | 0.609 | 0.635 | 0.637 | 0.605 | 0.676 | 0.658 | 0.570 |
| C>A | 0.858 | 0.744 | 0.904 | 0.802 | 0.894 | 0.900 | 0.893 | 0.859 | 0.908 | 0.857 |
| C>G | 0.757 | 0.655 | 0.869 | 0.750 | 0.844 | 0.830 | 0.770 | 0.884 | 0.872 | 0.786 |
| C>T | 0.887 | 0.819 | 0.916 | 0.833 | 0.901 | 0.895 | 0.893 | 0.889 | 0.919 | 0.891 |
| CpG_C>A | 0.745 | 0.662 | 0.755 | 0.681 | 0.663 | 0.713 | 0.730 | 0.617 | 0.737 | 0.664 |
| CpG_C>G | 0.687 | 0.517 | 0.696 | 0.557 | 0.531 | 0.559 | 0.660 | 0.577 | 0.676 | 0.554 |
| CpG_C>T | 0.956 | 0.928 | 0.966 | 0.930 | 0.944 | 0.947 | 0.957 | 0.928 | 0.969 | 0.936 |
| **Mean** | 0.772 | 0.684 | 0.818 | 0.729 | 0.773 | 0.782 | 0.780 | 0.779 | 0.816 | 0.754 |
| **Median** | 0.757 | 0.662 | 0.869 | 0.750 | 0.844 | 0.830 | 0.770 | 0.859 | 0.872 | 0.786 |

**Table S14**. Model performance between models that include or exclude non-neighboring shape interactions on test data. The "maximum achievable $R^2$" column were calculated by directly comparing test data with the training data. The "Aggarwala" columns describe performances of the current state of the art model. The remaining columns represent models that we fitted. Regarding abbreviations, "sh" and "sc" represent shape and nucleotide (sequence context) features, and the "neibr" suffix represents that only neighboring interactions were included (see Methods). The mean and median values were computed based on $R^2$ values of the nine individual models and were not computed for performance changes to avoid ambiguity.

| Mutation class | Maximum achievable $R^2$ | sh1 | sh2 | sh2neibr | sc2 | sc2neibr | sc3 | sc3neibr | sc4 | sc4neibr | sh1_sc2 | sh1_sc2neibr |
|---|---|---|---|---|---|---|---|---|---|---|---|---|
| A>C | 0.666 | 0.300 | 0.539 | 0.579 | 0.413 | 0.326 | 0.528 | 0.407 | 0.582 | 0.446 | 0.455 | 0.386 |
| A>G | 0.961 | 0.615 | 0.920 | 0.922 | 0.756 | 0.649 | 0.895 | 0.739 | 0.929 | 0.751 | 0.800 | 0.721 |
| A>T | 0.750 | 0.489 | 0.672 | 0.673 | 0.543 | 0.496 | 0.597 | 0.551 | 0.591 | 0.573 | 0.569 | 0.555 |
| C>A | 0.905 | 0.579 | 0.884 | 0.868 | 0.666 | 0.608 | 0.816 | 0.702 | 0.877 | 0.765 | 0.731 | 0.683 |
| C>G | 0.867 | 0.451 | 0.837 | 0.834 | 0.572 | 0.476 | 0.757 | 0.618 | 0.841 | 0.718 | 0.628 | 0.569 |
| C>T | 0.864 | 0.714 | 0.871 | 0.870 | 0.785 | 0.719 | 0.869 | 0.801 | 0.878 | 0.819 | 0.817 | 0.718 |
| CpG_C>A | 0.534 | 0.559 | 0.631 | 0.635 | 0.587 | 0.527 | 0.607 | 0.571 | 0.610 | 0.595 | 0.574 | 0.539 |
| CpG_C>G | 0.417 | 0.440 | 0.532 | 0.517 | 0.484 | 0.397 | 0.535 | 0.458 | 0.527 | 0.497 | 0.491 | 0.456 |
| CpG_C>T | 0.934 | 0.867 | 0.918 | 0.925 | 0.908 | 0.850 | 0.928 | 0.889 | 0.931 | 0.912 | 0.916 | 0.909 |
| **Mean** | 0.766 | 0.557 | 0.756 | 0.758 | 0.635 | 0.561 | 0.726 | 0.637 | 0.752 | 0.675 | 0.665 | 0.615 |
| **Median** | 0.864 | 0.559 | 0.837 | 0.834 | 0.587 | 0.527 | 0.757 | 0.618 | 0.841 | 0.718 | 0.628 | 0.569 |

| Mutation class | sh1_sc3 | sh1_sc3neibr | sh1_sc4 | sh1_sc4neibr | sh2neibr_sc2 | sh2neibr_sc2neibr | sh2neibr_sc3 | sh2neibr_sc3neibr | sh2neibr_sc4 | sh2neibr_sc4neibr |
|---|---|---|---|---|---|---|---|---|---|---|
| A>C | 0.533 | 0.448 | 0.586 | 0.502 | 0.569 | 0.574 | 0.543 | 0.591 | 0.592 | 0.555 |
| A>G | 0.893 | 0.803 | 0.930 | 0.854 | 0.916 | 0.914 | 0.924 | 0.913 | 0.931 | 0.914 |
| A>T | 0.603 | 0.592 | 0.610 | 0.610 | 0.653 | 0.653 | 0.616 | 0.675 | 0.631 | 0.590 |
| C>A | 0.845 | 0.743 | 0.878 | 0.801 | 0.872 | 0.875 | 0.875 | 0.851 | 0.882 | 0.850 |
| C>G | 0.758 | 0.664 | 0.842 | 0.754 | 0.824 | 0.816 | 0.768 | 0.846 | 0.844 | 0.787 |
| C>T | 0.871 | 0.816 | 0.878 | 0.825 | 0.874 | 0.870 | 0.876 | 0.868 | 0.882 | 0.870 |
| CpG_C>A | 0.618 | 0.605 | 0.609 | 0.616 | 0.603 | 0.632 | 0.615 | 0.566 | 0.611 | 0.606 |
| CpG_C>G | 0.529 | 0.466 | 0.526 | 0.495 | 0.493 | 0.497 | 0.530 | 0.506 | 0.524 | 0.490 |
| CpG_C>T | 0.930 | 0.915 | 0.932 | 0.917 | 0.927 | 0.926 | 0.930 | 0.914 | 0.934 | 0.922 |
| **Mean** | 0.731 | 0.673 | 0.755 | 0.708 | 0.748 | 0.751 | 0.742 | 0.748 | 0.759 | 0.732 |
| **Median** | 0.758 | 0.664 | 0.842 | 0.754 | 0.824 | 0.816 | 0.768 | 0.846 | 0.844 | 0.787 |
